# Multistage gradient-based whole-cortex modeling for joint inference of regional circuit heterogeneity and interregional structural connectivity

**DOI:** 10.64898/2026.11.16.767192

**Authors:** Boran Yang, Zhenyuan Jin, Xiaoyu Chen, Douglas Zhou, Songting Li

## Abstract

Current non-invasive neuroimaging methods cannot directly resolve regional circuit heterogeneity and interregional structural connectivity, limiting mechanistic understanding of brain structure-function relationships and accurate whole-brain modeling. Here we introduce StaGIC (Staged Gradient-based Inference of Circuitry), a method that jointly infers regional circuit heterogeneity and model-constrained structural connectivity (SC) from resting-state functional connectivity (FC) and functional connectivity dynamics (FCD). Ablation and synthetic-recovery experiments show that staged optimization improves fits to functional data and ground-truth parameter recovery, with inferred SC remaining highly consistent across retained solutions from independent optimization runs. In human data, freely optimized regional parameters exhibit consistent relationships with cortical hierarchy across parcellations, while SC inference substantially improves FC reconstruction. In macaque and marmoset data, inferred SC shows stronger correlation with tracer-derived projections than diffusion MRI estimates, particularly for long-range connections. As a proof of concept in computational psychiatry, group-level models of major depressive disorder (MDD) reproduce MDD-associated FC differences and generate exploratory hypotheses about hierarchical differentiation and interregional projection organization. StaGIC thus provides a differentiable framework for inferring multiscale cortical organization from non-invasive functional imaging.

---

Resting-state activity provides a widely accessible window into the large-scale organization of the brain. Across species, resting brains exhibit reproducible patterns of spontaneous activity [1–3], and static functional connectivity (FC) derived from these signals summarizes stable features of functional network organization [4–6]. Altered FC has been reported across multiple psychiatric disorders and transdiagnostic dimensions of psychopathology [7–11]. FC, however, remains an observational statistic: it describes covariation among brain regions but cannot determine which local circuit properties and long-range interactions generate the observed network organization. Constraining the latent structural configurations compatible with these observations is essential for moving from statistical descriptions of brain activity toward mechanistic and potentially individualized models.

Biophysically grounded whole-brain models offer a principled route from functional statistics to candidate circuit mechanisms [12]. By embedding neural mass or neural field models within empirically derived structural connectomes, these approaches have reproduced characteristic properties of resting-state fMRI signals and approximated static FC across multiple datasets and model formulations [13–18]. Subsequent developments have introduced cortical heterogeneity through macroscale gradients and used analytical FC approximations for computationally tractable fitting [16], incorporated functional connectivity dynamics (FCD) to constrain temporal fluctuations beyond static FC [17], and optimized anatomically constrained generative coupling to characterize condition- or state-dependent effective connectivity [19–21]. Machine-learning estimators and automatic-differentiation frameworks have expanded high-dimensional parameter estimation for whole-brain models [22, 23]; for direct gradient-based fitting, however, long stochastic simulations still create a difficult trade-off between temporal information and gradient stability. Jointly estimating region-specific circuit dynamics and uncertain inter-regional coupling nevertheless remains a high-dimensional inverse problem, and the anatomical content of inferred coupling has rarely been tested against measurements withheld from model fitting.

Structural connectivity (SC) constitutes a major source of uncertainty in this inverse problem. Human SC is commonly estimated using diffusion MRI tractography, which is non-invasive and broadly applicable but has recognized limitations, including high false-positive rates, unreliable reconstruction of long-range projections, and a restricted dynamic range of estimated connection strengths [24, 25]. Comparisons with invasive tract-tracing data in macaques have identified long-range pathways that are not reliably recovered by diffusion imaging [25, 26]. Tracer-derived projection weights span several orders of magnitude and reveal numerous weak, directed pathways that remain difficult to characterize with diffusion tractography [25, 27]. Rodent tracer datasets likewise reveal an ultra-dense cortical graph with highly heterogeneous, area-specific connectivity profiles [28]. Because structural coupling directly shapes the dynamics generated by biophysical models, errors in the connectome can propagate into fitted parameters and complicate their mechanistic interpretation [29]. Connectome uncertainty therefore remains a major bottleneck in large-scale circuit modeling [12]. Estimating coupling from functional dynamics may reduce dependence on tractography, but it also makes the inverse problem more underdetermined and places greater demands on optimization and validation.

Regional heterogeneity compounds this difficulty. The primate cortex is organized along a hierarchy that links laminar architecture, myelin content, gene expression, and functional specialization [30–33]. Neuroimaging studies have further revealed a spatial gradient of structure–function coupling: functional activity is more tightly constrained by structural wiring in unimodal sensory areas and progressively less so in transmodal association cortex [34, 35]. Neurotransmitter receptor densities also follow distinct topo-graphical gradients associated with macroscale functional organization and cortical rhythms [36]. These observations provide strong motivation for allowing local circuit properties to vary across regions. Many widely used whole-brain models nevertheless adopt shared regional parameters. Recent models describe regional heterogeneity using maps such as the principal gradient of resting-state functional connectivity (RSFC) and the ratio of T1-weighted to T2-weighted MRI signal (T1w/T2w) [16, 17]. This strategy provides effective regularization but restricts regional variation to a predefined low-dimensional subspace. Fully region-specific optimization offers greater flexibility [15], yet becomes difficult to stabilize when local parameters and inter-regional coupling are both trainable, because different parameter combinations can produce similar functional statistics. A scalable framework must therefore accommodate region-specific circuit dynamics and uncertain coupling while retaining sufficient dynamical constraints and allowing the inferred quantities to be independently evaluated.

Here we introduce StaGIC (Staged Gradient-based Inference of Circuitry), a gradient-based framework that represents a mean-field model (MFM) as a trainable recurrent dynamical system. StaGIC uses backpropagation through time (BPTT) to estimate freely varying regional heterogeneity parameters together with model-constrained inter-regional coupling from resting-state FC and FCD. Its central methodological component is a multistage optimization strategy that progressively expands the trainable parameter set and the dynamical objectives, mitigating the instability of end-to-end optimization over long stochastic trajectories. The coupling matrix is initialized based on interregional distance and subsequently refined using functional dynamics, reducing direct dependence on diffusion tractography. StaGIC thus provides a unified differentiable framework for fitting regional heterogeneity and inter-regional coupling while preserving the biophysical structure of the underlying circuit model.

We evaluate StaGIC across human and non-human primate datasets. In human data, StaGIC improves FC reconstruction while retaining FCD constraints, and the optimized regional parameters show consistent relationships with cortical hierarchy across parcellations. We further validate the multistage design through ablation and synthetic data recovery, assess the consistency of inferred SC across independent optimization runs, and benchmark functional reconstruction and connectome inference against established whole-cortex parameter optimization approaches, obtaining higher FC correlations in the evaluated benchmark settings. In macaque and marmoset data, the inferred coupling corresponds more closely to tracer-derived connectivity than do diffusion MRI estimates, providing an anatomical validation of our method. As a proof-of-concept application to major depressive disorder (MDD), group-specific models reproduce FC differences between control and MDD groups and provide exploratory latent contrasts in hierarchical organization and inferred coupling architecture.

## Results

### StaGIC: a multistage gradient-based optimization framework for mean-field models

Current optimization strategies for biophysical whole-brain models face a dimensionality–stability tradeoff. Increasing region-specific degrees of freedom while allowing inter-regional coupling to vary jointly creates a high-dimensional inverse problem that is difficult to optimize stably over long stochastic trajectories. Consequently, regional heterogeneity is often constrained to low-dimensional parameterizations and SC is commonly fixed during fitting. StaGIC addresses this coupled optimization problem by treating the biophysical model as a recurrent dynamical system and training it directly against empirical restingstate FC and FCD statistics (Fig. 1). FCD quantifies how FC patterns vary over time. We compute an FC matrix within each sliding time window and define each FCD entry as the Pearson correlation between the vectorized upper-triangular off-diagonal FC entries from two windows. High FCD values therefore indicate similar connectivity patterns across the corresponding windows [37].

**Fig 1.**
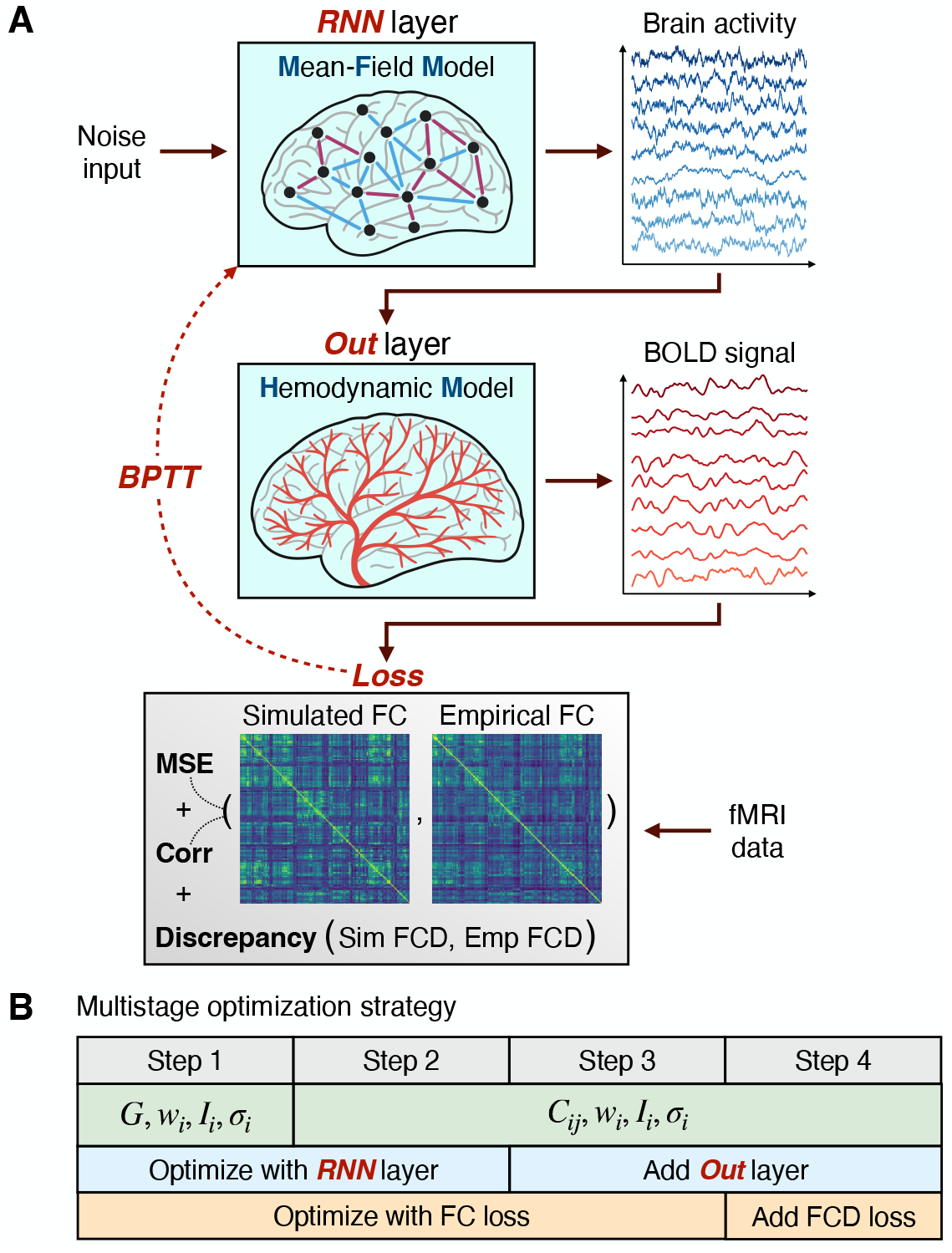
Schematic of StaGIC. (**A**) Noise-driven activity propagates through the MFM (“RNN layer”) and is transformed into BOLD signals by the hemodynamic output layer (“Outlayer”). Model parameters are optimized by BPTT using discrepancies between simulated and empirical FC and FCD. (**B**) Human optimization schedule. We first fit *G, w*_*i*_, *I*_*i*_, and *σ*_*i*_ to FC. We then fit *C*_*ij*_ along with the regional parameters, add the hemodynamic readout and introduce the FCD objective.

StaGIC uses BPTT within a multistage curriculum that progressively expands the trainable system. In the analysis of the human whole-cortex model illustrated in Fig. 1B, the optimization first allows the regional parameters (*w*_*i*_, *I*_*i*_, *σ*_*i*_) and the global coupling weight *G* to vary freely under an FC-based objective. After this initial regional and global parameter-optimization stage, the SC matrix *C*_*ij*_ is introduced for joint optimization. The hemodynamic model is then added as an output layer, followed by an FCD-based objective that constrains dynamic FC structure. During FCD-constrained training, BPTT minimizes discrepancies in the mean and standard deviation of sampled FCD entries. Training, validation and testing use the same empirical scans. In validation and test simulations, we draw new noise realizations and compare the distribution of FCD entries from full post-warm-up trajectories with the empirical distribution using a grid-based KS distance.

To test the contribution of the staged ordering, we compare the original StaGIC curriculum with three alternatives in which all parameters, all loss terms, or both are activated from the beginning. Across empirical and synthetic data, the original multistage strategy provides the strongest overall FC and FCD reconstruction, while the synthetic experiment further shows improved recovery of the groundtruth regional parameters and SC (Supplementary Fig. 1). The advantage is particularly pronounced for SC recovery, supporting progressive optimization as an effective strategy for constraining the coupled inverse problem.

Another key design choice in this curriculum is the initialization of the trainable SC matrix. The trainable matrix *C*_*ij*_ is initialized using the exponential distance rule (EDR) [38, 39], which defines a smooth distance-dependent connectivity scaffold from cortical geometry; the detailed construction is described in Methods. Diffusion MRI sets the decay length *λ* for each species. Given this scale, cortical geometry determines the initial weight of each connection. Functional fitting then updates the individual weights.

Each independent optimization run follows the complete multistage schedule and produces a sequence of saved parameter sets, termed candidate fits. For the run-level Glasser performance analysis in Fig. 3B, we select the candidate with the lowest validation loss among those saved after SC inference begins. This defines the run’s validation-selected fit; the one with the lowest validation loss across runs is the representative best fit.

**Fig 2.**
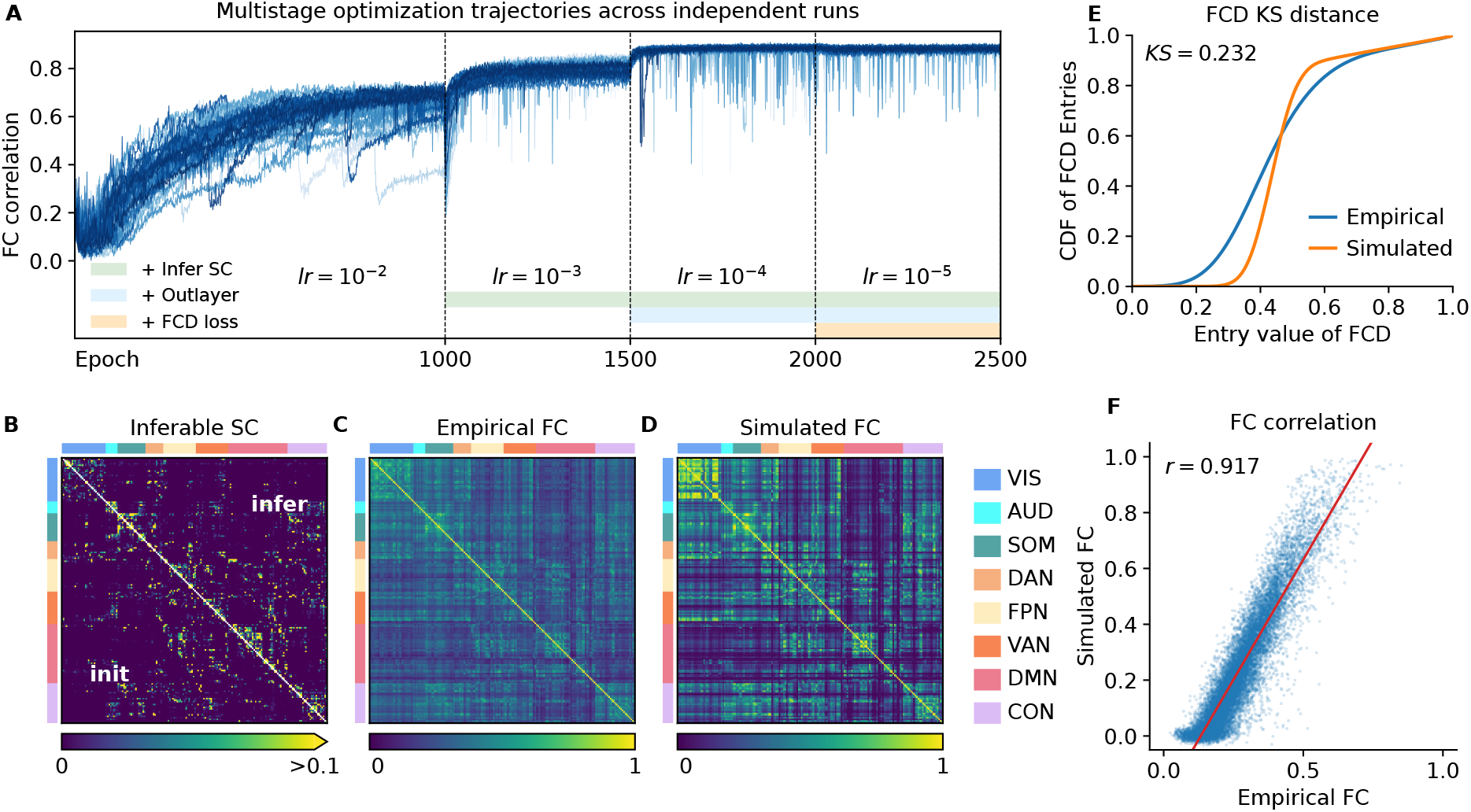
StaGIC accurately reconstructs static and dynamic resting-state organization. (**A**) Validation FC correlation across the Glasser multistage curriculum. Each curve traces the candidate fits sampled from one complete optimization run; dashed lines mark stage transitions, and horizontal bars indicate the introduction of SC inference, the hemodynamic output layer, and FCD loss. (**B**) SC from the representative best fit, with EDR-derived initial SC below and inferred SC above the diagonal. (**C, D**) Empirical FC and the elementwise mean of the 30 batch-specific simulated FC matrices from the representative best fit; ROIs in (**B**)–(**D**) follow the functional-network ordering shown at right. (**E**) Cumulative distributions of off-diagonal FCD entries. The empirical curve pools entries from all included resting-state fMRI runs, whereas the simulated curve pools entries from 30 independent test simulations of the representative best fit. The Kolmogorov–Smirnov (KS) distance is the maximum vertical separation between the two cumulative distributions. (**F**) Empirical versus simulated upper-triangular off-diagonal FC entries for the representative best fit; *r* denotes their Pearson correlation.

**Fig 3.**
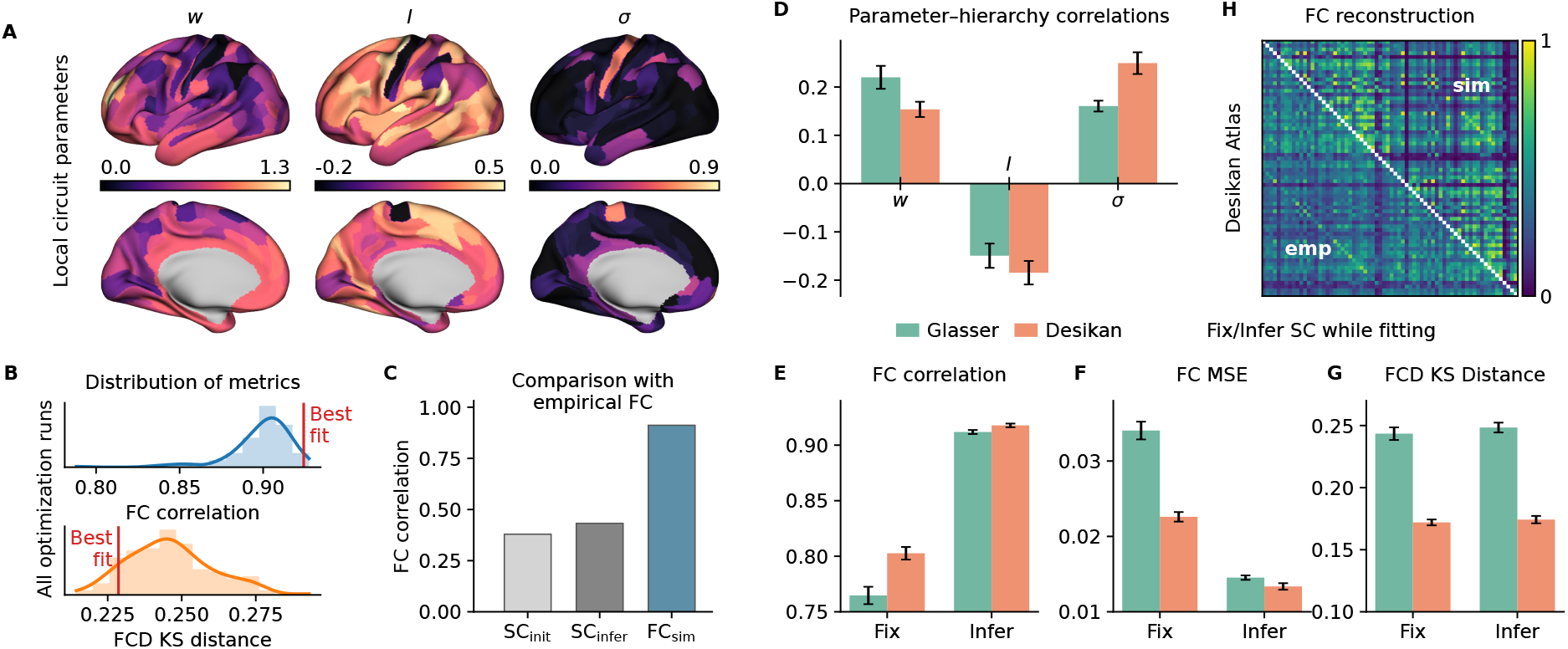
High-dimensional regional parameter inference preserves cortical hierarchy while improving functional reconstruction. (**A**) Glasser cortical maps of retained-fit mean *w*_*i*_, *I*_*i*_, and *σ*_*i*_, with ROI-level values assigned to the corresponding cortical parcels. (**B**) Test FC correlation and FCD KS distance, with one value per completed optimization run. For each run, the candidate fit with the lowest validation loss after SC inference begins is selected, without applying the ensemble-retention thresholds. Red lines mark the representative best fit used in Fig. 2B–F. (**C**) Correspondence of the EDR-derived initial SC, inferred SC, and simulated FC with empirical FC. Values show the mean across retained fits from the primary Glasser inferred-SC ensemble. (**D**) Parameter–hierarchy correlations across the Glasser and Desikan parcellations. (**E**)–(**G**) FC correlation, FC MSE, and FCD KS distance for models with EDR-derived SC held fixed and models with EDR-initialized trainable SC under both parcellations. (**H**) Desikan FC reconstruction for the representative best inferred-SC fit, with empirical FC below and simulated FC above the diagonal. Bars show means; error bars indicate 95% CI half-widths across retained fits in (**D**) and one validation-selected fit per completed optimization run in (**E**)–(**G**).

In the analysis of the human whole-cortex model, this multistage curriculum yields a progressive improvement in FC reconstruction under the Glasser parcellation. Validation FC correlation increases across the optimization stages, with marked gains after the introduction of SC inference and the hemodynamic output layer, and remains high during the final FCD-constrained stage (Fig. 2A; Supplementary Fig. 2). The inferred SC retains the broad geometry-dependent organization of the EDR initialization but contains structured deviations across functional networks that accompany the marked improvement in FC reconstruction (Fig. 2A,B).

The representative best fit closely reproduces the empirical FC matrix (Fig. 2C,D). Its simulated FCD-entry distribution also approximates the empirical distribution, yielding a KS distance of 0.232 (Fig. 2E).

The corresponding empirical and simulated FC entries show a Pearson correlation of *r* = 0.917 (Fig. 2F).

To assess SC consistency, we consider optimization runs that contribute at least one fit passing the prespecified validation screening described in Methods. From each of these runs, we use the retained fit with the lowest validation loss. The resulting SC matrices show strong agreement across independent optimization runs (Supplementary Fig. 3).

### High-dimensional regional parameter inference preserves cortical hierarchy while improving functional reconstruction

Freely optimizing regional circuit parameters tests whether improved functional reconstruction is accompanied by interpretable cortical organization. Previous models constrain regional heterogeneity using predefined cortical gradients [16, 17]. For example, the parametric mean-field model (pMFM) expresses *w*_*i*_, *I*_*i*_, and *σ*_*i*_ as linear combinations of T1w/T2w and the principal RSFC gradient. StaGIC instead estimates these parameters at ROI resolution without imposing those spatial profiles. We therefore examine whether cortical hierarchy remains evident after optimization.

The optimized parameters form coherent cortical patterns under the Glasser parcellation (Fig. 3A). Functional performance is also consistent across completed optimization runs: FC correlations remain high and FCD KS distances occupy a relatively narrow range when each run contributes its validation-selected fit (Fig. 3B).

Direct comparison with empirical FC shows that SC inference only modestly increases SC–FC similarity, whereas the recurrent model produces much stronger agreement between simulated and empirical FC (Fig. 3C).

In both parcellations, we compared the freely estimated parameter maps with the principal gradient of the empirical FC target. Recurrent strength (*w*_*i*_) and noise amplitude (*σ*_*i*_) correlated positively with this gradient, and external input (*I*_*i*_) correlated negatively (Fig. 3D). Recurrent strength also increased toward association cortex in the best-fitting pMFM solution [17]. An earlier model found the strongest recurrence in sensorimotor cortex [15].

Compared with models in which EDR-derived SC is held fixed, SC inference increases FC correlation and reduces FC MSE, with broadly similar FCD KS distances under both parcellations (Fig. 3E–G). Strong FC reconstruction is also obtained under the Desikan parcellation (Fig. 3H).

### Comparisons with tracer-derived SC of non-human primates support anatomical correspondence of inferred SC

An independent anatomical test of StaGIC-inferred SC is its correspondence with tracer-derived projections. The human analyses show that StaGIC infers a latent SC that enhances resting-state reconstruction while preserving interpretable cortical organization. Yet anatomical validation in humans remains limited because diffusion-based SC provides only an indirect and error-prone approximation of inter-regional projections. To move beyond this limitation, we apply StaGIC to macaque and marmoset resting-state fMRI, where tracer-derived connectivity provides an invasive anatomical reference for inter-regional projections (Fig. 4).

**Fig 4.**
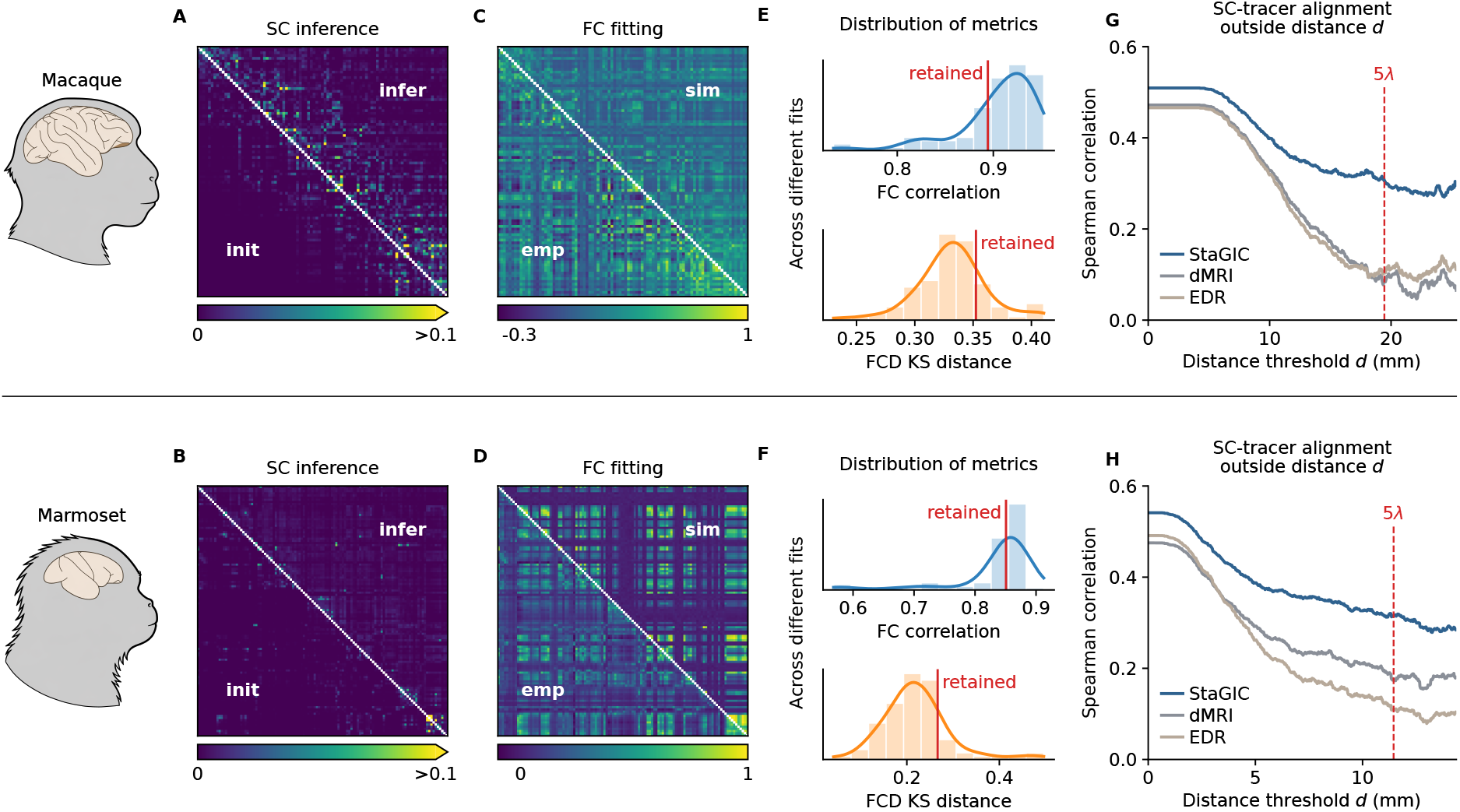
Connectome inference and tracer validation in non-human primates. Macaque results are shown above and marmoset results below. (**A, B**) Normalized SC matrices, with EDR-derived initial SC below and retained-fit mean inferred SC above the diagonal. (**C, D**) Empirical FC below and retained-fit mean simulated FC above the diagonal. (**E, F**) Test FC-correlation and FCD-KS-distance distributions across the 20 lowest-validation-loss eligible candidate fits from each optimization run, including both retained and non-retained shortlisted fits; red lines mark, respectively, the minimum test FC correlation and maximum test FCD KS distance attained among retained fits. The red lines are post hoc descriptive references and are not used for fit selection. (**G, H**) Spearman correlations of dMRI-derived, EDR-derived, and StaGIC-inferred SC with the fraction of labeled neurons (FLN) across minimum distance thresholds. The analysis includes measured zero values. Candidate SC matrices are symmetrized, whereas FLN retains its directed, target-normalized representation. The StaGIC curves are computed by averaging SC across retained fits before correlation; red dashed lines mark the species-specific 5*λ* threshold.

This analysis of non-human primate datasets evaluates both the generalizability of StaGIC and the anatomical correspondence of its inferred connectome. Macaque and marmoset datasets differ in parcellation, acquisition duration, distance-decay structure, and tracer coverage, yet both allow model-inferred SC to be compared with invasive projection measurements. In both species, the inferred SC also departs from the EDR-derived initialization, indicating that optimization does not simply rescale the distance prior but reshapes the latent structural substrate that supports the observed dynamics (Fig. 4A,B). Despite these differences, StaGIC reproduces species-specific FC organization in both primates, with high test FC correlations and low KS distances of FCD across different fits (Fig. 4C–F).

We then test whether the optimized SC contains anatomical information beyond diffusion-based estimates and the EDR initialization. For each species, we compare dMRI-derived SC, EDR-derived initial SC, and StaGIC-inferred SC with tracer-derived connectivity across anatomical distance thresholds. Here, *λ* denotes the species-specific characteristic decay length of the EDR [38, 39]. We classify a region pair as long-range when its separation exceeds 5*λ*. At each threshold *d*, tracer-covered ROI pairs with anatomical distance greater than *d* are retained, and Spearman’s rank correlation is computed between tracer connectivity and each SC estimate while preserving measured zero-valued tracer entries. In both species, the symmetrized fitted weights correlated more strongly with directed tracer measurements than did either the EDR weights or the tractography streamline counts (Fig. 4G,H). The difference persisted when only pairs separated by more than 5*λ* were included.

### StaGIC extends to group-level modeling of MDD functional alterations

MDD provides a proof-of-concept setting for applying StaGIC to disease-associated functional alterations. Distributed resting-state FC alterations are well documented in MDD [11, 40], whereas their underlying circuit configurations remain less directly accessible. We fit separate group-level models to the group-averaged FC and pooled FCD distributions of normal controls (NC) and patients with MDD from REST-meta-MDD [11]. Both models use the 96-region Harvard–Oxford parcellation provided with the released dataset and the same multistage optimization schedule.

The fitted models reproduce the empirical FC targets in both NC and MDD (Fig. 5A,B). To compare the two conditions, we compute the elementwise difference ΔFC = FC^NC^ − FC^MDD^ for both empirical and simulated FC matrices. Both contrast matrices show predominantly positive differences, indicating stronger FC in NC across much of the cortex (Fig. 5C,D). FC correlations remain high across different fits in both groups (Fig. 5E).

**Fig 5.**
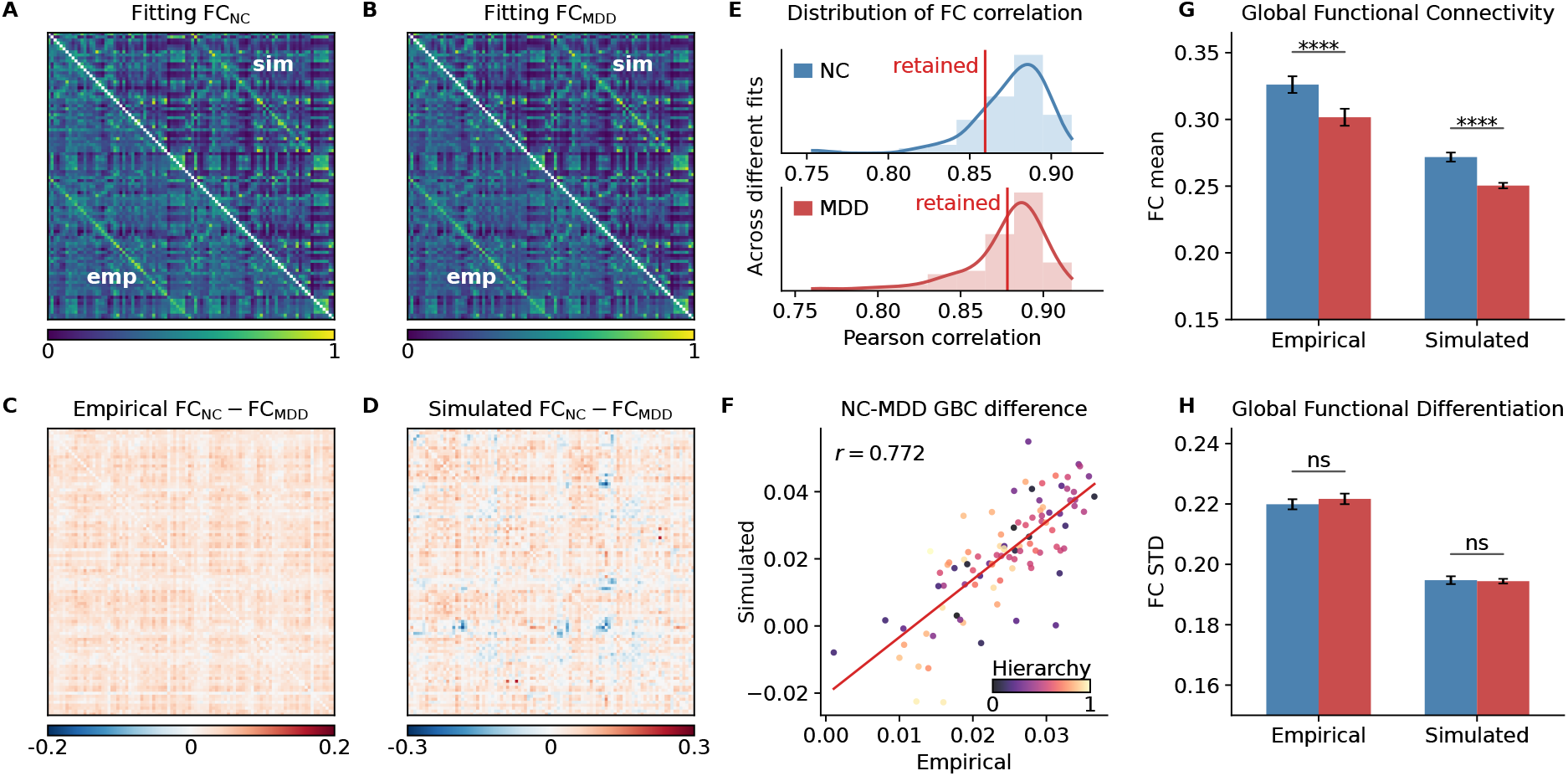
Group-level models reconstruct MDD-associated functional alterations. (**A, B**) NC and MDD FC fits, with empirical FC below and retained-fit mean simulated FC above the diagonal. (**C, D**) Empirical and simulated NC–MDD FC contrasts in the same ROI order; positive values indicate stronger FC in NC. (**E**) Test FC-correlation distributions across the 20 lowest-validation-loss eligible candidate fits from each optimization run in NC and MDD, including both retained and non-retained shortlisted fits; red lines mark the group-specific minimum test FC correlation attained among retained fits. The red lines are post hoc descriptive references and are not used for fit selection. (**F**) Empirical versus simulated NC– MDD differences in ROI-level GBC, defined as each ROI’s mean FC with all other ROIs; points are colored by the shared NC–MDD reference hierarchy. (**G, H**) Group differences in global functional connectivity and differentiation, defined as the mean and standard deviation of off-diagonal FC entries, respectively. Bars show means; error bars indicate 95% CI half-widths across participants for empirical data and retained fits for simulated data.

This correspondence extends from the full FC matrix to regional and global network summaries. At the ROI level, global brain connectivity (GBC) is defined as the mean FC between an ROI and all other ROIs. The empirical and simulated NC–MDD GBC differences are correlated across ROIs (*r* = 0.772; Fig. 5F), indicating that the models capture regional variation in the magnitude of the observed FC differences.

The models also reproduce the direction of the global FC alteration: global functional connectivity, defined as the mean off-diagonal FC, is lower in MDD than in NC in both empirical and simulated data, whereas global functional differentiation, defined as the standard deviation of off-diagonal FC entries, shows no significant group difference in either case (Fig. 5G,H).

### Latent parameter inference provides circuit-level hypotheses for MDD-associated functional differences

We next compare the inferred regional parameters and interregional projections between the NC and MDD models.

Using a shared reference hierarchy derived from the mean of the empirical NC and MDD FC matrices, we find lower absolute parameter–hierarchy correlations for *w*_*i*_, *I*_*i*_, and *σ*_*i*_ in the MDD model (Fig. 6A,D). This pattern is consistent with reduced hierarchical differentiation of the inferred regional circuit properties.

**Fig 6.**
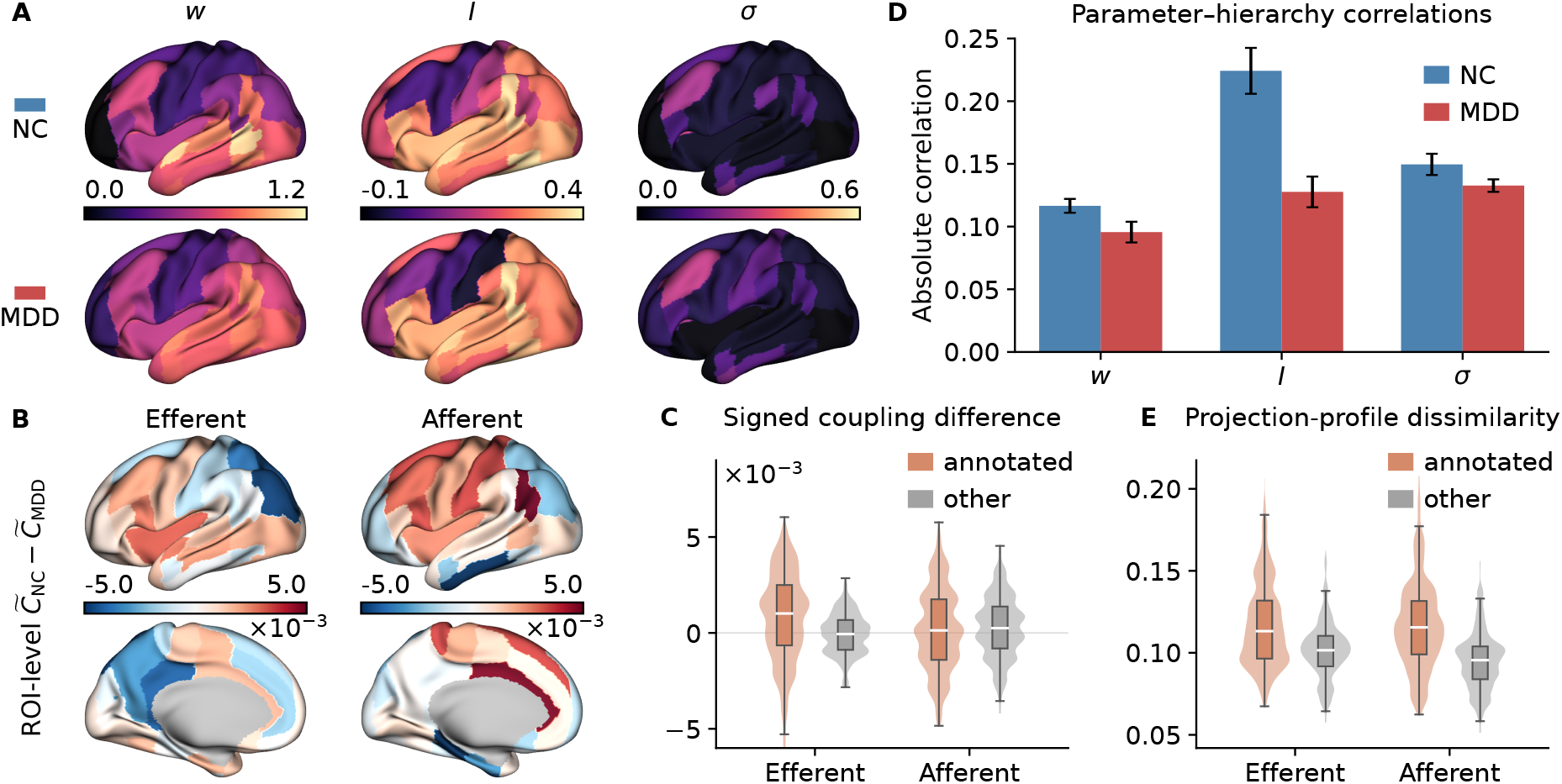
MDD group models yield exploratory contrasts in cortical hierarchy and projection architecture. (**A**) NC and MDD maps of retained-fit mean *w*_*i*_, *I*_*i*_, and *σ*_*i*_, shown with shared group-wise color ranges. (**B**) Mean signed NC–MDD differences in effective inferred coupling, 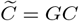, across all cross-fit pairs; efferent and afferent maps summarize source- and target-ROI differences, respectively. (**C**) Cross-fit distributions of category-averaged signed differences in effective inferred coupling for MDD-annotated and other ROIs; positive values indicate stronger coupling in NC. (**D**) Absolute correlations of regional parameters with the shared Harvard–Oxford reference hierarchy in NC and MDD. (**E**) Cross-fit projection-profile dissimilarity, defined as one minus the NC–MDD Pearson correlation, for efferent and afferent profiles of MDD-annotated and other ROIs. In (**C**) and (**E**), violin widths indicate distribution density, and embedded boxes show medians and interquartile ranges. Bars in (**D**) show means *±* 95% CI half-widths across retained fits.

The inferred SC provides a complementary view of inter-areal projection architecture. To contextualize these projection differences with prior MDD literature, selected Harvard–Oxford ROIs are designated MDD-annotated through a literature-informed mapping, with all remaining cortical ROIs forming the comparison set (Supplementary Table 1) [40–45]. Across pairs of fitted models, the median NC–MDD difference in mean outgoing coupling was positive for MDD-annotated regions and close to zero for other regions. Median differences in incoming coupling were close to zero in both sets (Fig. 6B,C). The annotated regions also had higher median dissimilarity between NC and MDD coupling profiles in both directions (Fig. 6E).

## Discussion

StaGIC addresses a coupled inverse problem in whole-brain modeling by jointly inferring regional circuit heterogeneity and latent inter-regional coupling from spontaneous functional dynamics. Its central contribution is to combine freely varying regional parameters, trainable SC, biophysical recurrent dynamics, and FC/FCD constraints within a multistage gradient-based framework. In human data, SC inference substantially improves functional reconstruction while the optimized regional parameters preserve a consistent relationship with cortical hierarchy across parcellations. Ablation and synthetic-recovery analyses show that the multistage curriculum improves both functional fitting and ground-truth parameter recovery, while inferred SC remains highly consistent across independent optimization runs. Macaque and marmoset tracer validation further connects the inferred SC to independently measured anatomical projections.

The MDD application links group-level functional alterations to exploratory hypotheses of attenuated hierarchical differentiation and preferential efferent coupling changes involving MDD-annotated regions.

StaGIC retains interpretable cortical organization despite its high-dimensional parameterization. Regional parameters are allowed to vary freely across ROIs without imposing a predefined cortical gradient, yet their relationship with cortical hierarchy remains consistent across parcellations. Similarly, the inferred SC retains the broad geometry-dependent organization of its initialization yet produces large improvements in FC reconstruction, indicating that structured changes in coupling can have substantial consequences after recurrent dynamical transformation. These observations motivate a view of whole-brain fitting in which biological organization emerges from the interaction between anatomical priors, functional constraints, and biophysical dynamics.

StaGIC builds on recent efforts to incorporate regional heterogeneity and automatic differentiation into whole-brain model fitting. In the benchmarks evaluated here, it achieves higher FC correlations than the comparison methods. pMFM introduces biologically informed heterogeneity by mapping anatomical and functional cortical gradients onto regional circuit parameters [17], providing effective low-dimensional regularization. StaGIC instead permits ROI-specific values of *w*_*i*_, *I*_*i*_, and *σ*_*i*_ within a jointly optimized recurrent system and evaluates whether hierarchical cortical organization emerges after fitting. In fixed-SC benchmarks, StaGIC achieves higher FC correlations than pMFM under both Desikan and Glasser parcellations (Supplementary Fig. 4A).

TVB-Optim independently establishes automatic differentiation as an efficient strategy for high-dimensional whole-brain parameter estimation [23]. Under trainable-SC settings, StaGIC achieves higher FC correlations than TVB-Optim under both human parcellations, with a particularly pronounced advantage under the Glasser parcellation (Supplementary Fig. 4C). The advantage extends to macaque and marmoset, where StaGIC achieves higher FC correlations and stronger SC–tracer rank correspondence across most evaluated distance thresholds (Supplementary Fig. 4E,G,H). The tracer analyses complement these functional benchmarks with an independent anatomical evaluation of inferred SC.

The multistage curriculum is an important component of the connectome inference strategy. The ablation analysis compares the original StaGIC schedule with alternatives in which all parameters, all losses, or both are activated from the beginning. Across empirical and synthetic datasets, the original curriculum provides the best overall FC and FCD reconstruction, while synthetic experiments further show substantially improved recovery of the ground-truth regional parameters and SC (Supplementary Fig. 1). The advantage is particularly clear for SC recovery, indicating that optimizing regional and global parameters before introducing trainable SC and later dynamical objectives helps constrain an otherwise highly coupled inverse problem.

In the Glasser analysis, we compared one coupling matrix per retained run, selecting the retained fit with the lowest validation loss in each. These matrices were similar across runs that began from the same EDR initialization (Supplementary Fig. 3). The primate tracer data provide a separate anatomical test. In both species, the symmetric component of fitted coupling had a higher rank correlation with tracer projection strengths than did the initial EDR weights or the tractography streamline counts (Fig. 4G,H). Each correlation pools region pairs across the range of distances above its threshold.

Different combinations of regional parameters and directed coupling weights can produce similar FC and FCD. Training uses short trajectories and constrains the mean and standard deviation of sampled FCD entries. For each fit, averaging FC matrices and pooling FCD entries across 30 new simulations reduces variation due to simulation noise. Averaging parameter maps and coupling matrices across retained fits summarizes the solutions obtained for the same empirical target.

Extending StaGIC to task-evoked fMRI could connect the intrinsic dynamical organization identified at rest with temporally structured cognition. Task timing and evoked BOLD trajectories provide explicit constraints on how the resting-state-derived connectome responds to external inputs and supports context-dependent computation. Such models may help distinguish stable circuit properties from transient regional or inter-areal changes recruited by specific cognitive demands, providing a route from the intrinsic dynamical substrate of the brain toward mechanistic models of cognition and conscious processing.

A second direction is the development of scalable subject-specific models. A practical strategy is to first infer a robust group-level model and then fine-tune a restricted subset of parameters for each individual, using the group solution as a biologically grounded initialization. Determining which parameters should be individualized remains an important scientific question, as regional circuit properties, selected SC projections, global coupling, and noise amplitudes may encode distinct sources of individual or disease-related variability. This transition also requires substantial computational acceleration, because reliable subject-level inference will require multiple fits for each individual and repeated optimization across large cohorts to characterize solution uncertainty. Faster simulation and gradient computation, memory-efficient BPTT, and parallelized training and validation could make this workflow practical while preserving the mechanistic interpretability of StaGIC. A first test of an individual model is whether parameters fitted to one scan predict FC and FCD in a later scan of the same person. In depression, longitudinal data could also test whether changes in fitted coupling track changes in symptoms.

## Online Methods

### Data

We evaluate StaGIC using resting-state fMRI datasets from healthy humans, macaques, marmosets, and patients with MDD. All analyses use previously collected data and involve no new human or animal data acquisition. Ethics approvals and informed-consent procedures for the human datasets, and animal-care approvals for the non-human primate datasets, are reported in the corresponding source publications [11, 27, 46–49].

The cross-species analyses use hemispheric cortical representations within a common EDR-based initialization and model-fitting framework. For the human analysis, we use the Glasser parcellation with 180 cortical regions per hemisphere [50], following the within-hemisphere modeling scope of Demirtaş et al. [16]. After run-level quality control, we retain 4,227 usable MSMAll 3-T resting-state runs from 1,090 participants from the WU–Minn Human Connectome Project [46]. For each run, Pearson FC is computed, and the left- and right-hemisphere within-hemisphere FC blocks are aligned by homologous parcels and averaged elementwise. This yields a 180×180 FC matrix for each run without cross-hemispheric edges. The empirical target is the elementwise arithmetic mean of these hemisphere-averaged run-level FC matrices on the native *r* scale without Fisher *z* transformation.

After dataset-specific quality control, the macaque analysis retains 32 resting-state runs from eight animals from the dataset of Shen et al. [47]. Cortical time series are represented in the 91-area Markov hemispheric parcellation, and tracer-derived FLN is obtained from the corresponding weighted directed interareal matrix [27].

The marmoset analysis uses 710 valid resting-state runs from 113 sessions in 39 animals from Tian et al. [48], represented in the 116-area MBMv3/Paxinos hemispheric parcellation [49, 51]. Marmoset tracer connectivity is obtained from the open cellular-resolution resource of Majka et al. [49]. The published marmoset tracer matrix contains 3,474 nonzero FLN entries [52]. For both non-human primate datasets, empirical FC targets are the elementwise means of the corresponding run-level Pearson FC matrices. Neither tracer dataset is used during model fitting.

For bilateral cortical modeling, we analyze the same HCP runs using all 68 regions of the Desikan– Killiany parcellation [53]. This provides an additional human parcellation and a common bilateral setting for comparisons with pMFM [17] and TVB-Optim [23]. Run-level Pearson FC matrices retain both within- and cross-hemispheric edges, and their elementwise arithmetic mean on the native *r* scale defines the 68×68 empirical target.

The MDD analysis uses the publicly released Harvard–Oxford parcel-wise data from the REST-meta-MDD project (https://rfmri.org/REST-meta-MDD), derived from the consortium described by Yan et al. [11]; all 96 released ROIs across both hemispheres are retained. After excluding scans with fewer than 190 volumes, 968 NC and 1,150 MDD participants remain, each contributing one scan. Participant-level Pearson FC matrices are averaged elementwise within each group to construct the NC and MDD fitting targets. The empirical summaries in Fig. 5G,H are instead computed separately for each participant.

FCD is computed by sliding a temporal window across each regional time series, vectorizing the upper-triangular off-diagonal FC entries in each window, and correlating the resulting window-wise FC vectors [37]. For each dataset or diagnostic group, off-diagonal FCD entries are pooled directly across all included empirical runs or participants, and a single empirical cumulative distribution is then constructed from the pooled entries. Participant- or run-specific cumulative distributions are not averaged.

### Construction of the initial connectome

We construct the initial connectome from vertex- or voxel-level spatial geometry using an exponential distance rule (EDR), with the decay constant *λ* calibrated separately for each species as described below. The procedure starts from two atlas files: a pairwise Euclidean distance matrix across surface vertices or voxels, and a label vector that assigns each vertex or voxel to one atlas region. Let *d*_*uv*_ denote the distance between elements *u* and *v*, and let *λ* denote the species-specific distance-decay constant. Here, *λ* denotes the characteristic EDR decay length.

The target connection-length density is modeled using the exponential distance rule (EDR) [38, 39]:

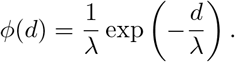

Under this exponential length model, the probability mass beyond 5*λ* is *e*^*−*5^ ≈ 0.0067. We therefore use 5*λ* as an operational threshold for examining connections in the long-distance tail.

Only the lower triangle of the vertex/voxel distance matrix is used to estimate the distance distribution. Distances are binned at 0.1-mm resolution. For a distance bin *b* with center *c*_*b*_, width Δ*d* = 0.1 mm, and *n*_*b*_ element pairs, the bin-specific per-pair connection probability is

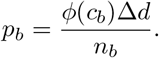

For numerical implementation, zero bin counts are temporarily replaced by one before division; because empty bins contain no element pairs, this safeguard assigns neither connections nor probability mass to those bins. Each off-diagonal element pair (*u, v*) is then assigned the probability *p*_*b*(*u,v*)_ corresponding to the bin containing *d*_*uv*_, producing a symmetric vertex/voxel-level probability matrix *P* .

The vertex/voxel probabilities are converted to an ROI-level initial SC by matrix aggregation through the atlas labels. We construct a binary membership matrix *M*, where *M*_*ur*_ = 1 if element *u* belongs to region *r* and *M*_*ur*_ = 0 otherwise. The EDR basis connectome is then computed as

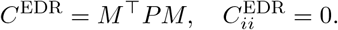

Thus, each ROI-level entry is the sum of EDR probabilities across all vertex or voxel pairs linking the two labeled regions. The resulting matrix serves as the initial structural substrate before connectome inference.

The decay constant *λ* sets the single global spatial scale of the EDR prior and is calibrated separately for each species. Diffusion MRI data were obtained from the WU–Minn HCP for the human analyses [46], the macaque connectome resource of Shen et al. [47], and the integrated marmoset resource of Tian et al. [48]. Diffusion tractography was performed using DIPY [54], with fiber orientations reconstructed by constrained spherical deconvolution (CSD) and local streamline tracking terminated using an FA threshold of 0.2. SC was constructed by counting streamlines whose endpoints fell within each pair of cortical regions. The resulting matrices were symmetrized, their diagonal elements were set to zero, and edge weights were normalized by the maximum off-diagonal streamline count. We determine *λ* by grid search against diffusion-derived SC: for each candidate value, an ROI-level EDR connectome is generated from cortical geometry as described above, and the Pearson correlation between its off-diagonal entries and the corresponding diffusion-derived SC is evaluated. The value maximizing this correspondence is selected before any functional-model optimization or tracer-based evaluation. This calibration determines only the characteristic distance-decay scale; individual EDR connections remain defined by cortical geometry and do not inherit tractography-derived edge topology or weights. Human analyses search *λ* from 5 to 15 mm, macaque analyses from 1 to 10 mm, and marmoset analyses from 1 to 5 mm, in 0.01-mm increments. This procedure yields *λ* = 6.99 mm for human cortical analyses, *λ* = 3.89 mm for macaque, and *λ* = 2.29 mm for marmoset.

### Dynamic mean-field model

The mean-field model (MFM) is obtained by applying a mean-field reduction to a detailed spiking neuronal network [13]. In its original formulation, the model uses the Abbott–Chance transfer function and is described by the following system of nonlinear stochastic differential equations:

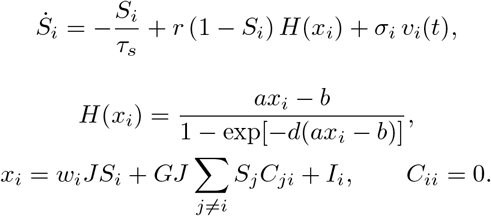

Here, *x*_*i*_, *H*(*x*_*i*_), and *S*_*i*_ denote the total input current, the population firing rate, and the average synaptic gating variable of the *i*-th cortical region, respectively. The total input current *x*_*i*_ is jointly determined by the local recurrent connection strength *w*_*i*_, the excitatory input *I*_*i*_, and long-range inter-regional interactions. The structural connectivity *C*_*ij*_ specifies the directed coupling strength from source region *i* to target region *j*, thereby modulating inter-regional information flow. The global coupling parameter *G* scales the contribution of long-range inputs relative to local recurrent and external inputs.

Throughout this study, SC denotes only the off-diagonal inter-regional entries *C*_*ij*_ with *i≠ j*, and all reported SC matrices have *C*_*ii*_ = 0. Local recurrent strengths *w*_*i*_ are directly optimized regional parameters. During SC-inference stages, *w*_*i*_, *I*_*i*_, and *σ*_*i*_ remain trainable alongside inter-regional SC.

Following previous work [13], the synaptic coupling is fixed at *J* = 0.2609 nA. The parameters for the input–output function *H*(*x*_*i*_) are set to *a* = 270 (n/C), *b* = 108 (Hz), and *d* = 0.154 (s). The kinetic parameters governing synaptic activity are *r* = 0.641 and *τ*_*s*_ = 0.1 s. The noise term *v*_*i*_(*t*) represents temporally uncorrelated Gaussian white noise scaled by the region-specific amplitude *σ*_*i*_.

The MFM is a reduced population-level description that retains interpretable variables such as firing rate and synaptic gating while using far fewer state variables than the detailed spiking-network description from which it is derived [13].

In the main StaGIC pipeline, the Abbott–Chance transfer function is replaced by a smooth Softplus approximation,

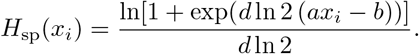

This parameterization matches both the value *H*(*b/a*) = 1*/d* and the local slope *H*^*′*^(*b/a*) = *a/*2 of the original transfer function. The approximation is evaluated using a numerically stable log-sum-exp operation and is used for training, validation and test simulations of the main model.

### Hemodynamic model

The main optimization pipeline transforms simulated neural activity into BOLD-like signals using the two-state linear hemodynamic filter defined below, which we refer to as the Volterra readout. We use this readout because its low-dimensional state-space form is computationally efficient for BPTT while preserving the FC and FCD statistics needed for model fitting. This layer is implemented as a two-state linear system,

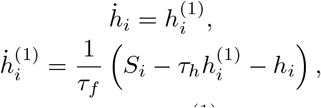

where *h*_*i*_ is the simulated BOLD response of region *i*, 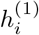 is the auxiliary derivative state, and (*τ*_*h*_, *τ*_*f*_) = (2.3s, 3.9s^2^). The Volterra output *h*_*i*_ is used in the later optimization stages, validation simulations, test simulations, and the quantitative FC/FCD results reported for the main model.

To assess the combined effect of the neural and hemodynamic approximations, we compare the main implementation, which combines the Softplus transfer function with the Volterra readout, with a reference implementation using the original Abbott–Chance transfer function and the Balloon–Windkessel hemo-dynamic model [55]. Both implementations use the same optimized neural parameters. The resulting FC and FCD summaries are highly consistent across the Glasser and Desikan atlases (Supplementary Fig. 5). In the Balloon–Windkessel model, synaptic activity *S*_*i*_ drives changes in a vasodilatory signal *z*_*i*_, which subsequently modulates blood inflow *f*_*i*_, blood volume *v*_*i*_, and deoxyhemoglobin content *q*_*i*_ according to:

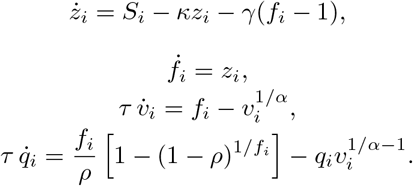

The resting oxygen extraction fraction is fixed at *ρ* = 0.34. The kinetic parameters include a signal decay rate *κ* = 0.65 s^*−*1^, a flow-dependent elimination rate *γ* = 0.41 s^*−*1^, a hemodynamic transit time *τ* = 0.98 s, and Grubb’s exponent *α* = 0.32, consistent with prior work [13]. The BOLD signal for region *i* is computed as [56, 57]:

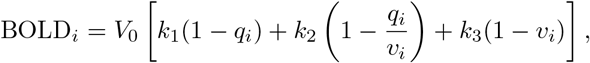

where *V*_0_ = 0.02 denotes the resting blood volume fraction. The coefficients (*k*_1_, *k*_2_, *k*_3_) depend on magnetic-field and acquisition parameters [57]:

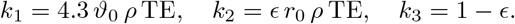

Here, *ϑ*_0_ = 28.265*B*_0_ is the frequency offset at the surface of magnetized vessels, with a main magnetic field strength of *B*_0_ = 3 T for the HCP dataset. At 3 T, the intravascular relaxation rate is *r*_0_ = 110 Hz and the intravascular–extravascular signal ratio is *ϵ* = 0.47 [15, 56, 57]. The HCP echo time is TE = 33.1 ms [15, 46].

Both the MFM and the hemodynamic model are integrated using Euler’s method with a 1 ms time step. The initial values of *S*_*i*_ are randomly initialized. Validation and test simulations begin with a 2 min warm-up period that is discarded before metric computation. The analyzed post-warm-up simulation duration is 14.4 min for the HCP human analyses, 380 s for the Harvard–Oxford analyses, 1200 s for macaque, and 1024 s for marmoset. The synthetic BOLD signals are downsampled to 0.72 s or 2.00 s— depending on species and dataset—to match the temporal resolution of the corresponding empirical fMRI data.

### Gradient-based estimation of model parameters

StaGIC implements the MFM as a recurrent dynamical system and estimates model parameters using backpropagation through time (BPTT). The trainable variables across the multistage curriculum include regional recurrent strengths *w*_*i*_, external inputs *I*_*i*_, regional noise amplitudes *σ*_*i*_, the global coupling parameter *G*, and, when connectome inference is active, a full unconstrained raw coupling matrix Θ. The off-diagonal entries of the raw coupling matrix are mapped to nonnegative inter-regional weights as

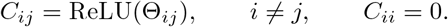

Local recurrent strengths *w*_*i*_ remain separate, directly optimized regional parameters during SC inference. No symmetry constraint is imposed on the off-diagonal entries, so Θ_*ij*_ and Θ_*ji*_ are independently trainable. All reported SC quantities are derived from the resulting nonnegative off-diagonal *C*_*ij*_ rather than from the raw optimization variable Θ_*ij*_; analyses explicitly described as effective coupling use 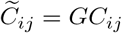.

At each training epoch, stochastic inputs are sampled as Gaussian white noise and scaled by the regional noise amplitudes. The model is first simulated through a warm-up period and is then unrolled over the training interval for loss evaluation. The FC-based objective compares the simulated and empirical FC matrices using both correlation and mean-squared error:

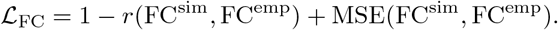

For optimization stages that include FCD constraints, an additional loss term is used to match the distributional summary of simulated and empirical FCD entries:

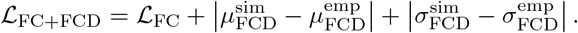

Here, *µ*_FCD_ and *σ*_FCD_ denote the mean and standard deviation of sampled FCD entries, respectively.

Parameters are optimized with Adam. To improve numerical stability, gradients are clipped element-wise to [−10^5^, 10^5^]. If an individual trainable parameter becomes NaN while the loss remains finite, the affected entry is reset to its value from the preceding epoch plus 10^*−*5^(*U* + 1), where *U ∼* Uniform(0, 1). Optimization runs producing a NaN loss are terminated and excluded from downstream analyses.

### Multistage optimization procedure

StaGIC uses a multistage optimization curriculum that gradually increases model complexity, comprising four optimization components: regional/global parameter fitting, SC inference, hemodynamic-output fitting, and FCD refinement. The relative ordering of SC inference and hemodynamic fitting is adjusted across datasets. This curriculum allows shorter and simpler simulations to stabilize the dynamical system before longer BOLD-level and FCD-constrained optimization.

In the initial stage, the model optimizes regional heterogeneity and global coupling using neural activity-level outputs and the FC objective. When connectome inference begins, *G* is held at its fitted value and the off-diagonal weights *C*_*ij*_ are optimized jointly with *w*_*i*_, *I*_*i*_, and *σ*_*i*_ (Supplementary Table 2). The hemodynamic output layer transforms simulated neural activity into BOLD-like signals for comparison with empirical FC. In the final stage, the combined FC+FCD objective is used to refine the inferred parameters under both static and dynamic functional constraints.

Minor dataset-specific adjustments are made to accommodate differences in parcellation, sampling rate, empirical scan duration, and the relative ordering of SC inference and hemodynamic-output fitting. For example, in the human analyses, SC inference is introduced before the hemodynamic output layer, whereas in the non-human primate analyses, the hemodynamic output layer is introduced before SC inference. Despite these differences, all optimization schedules comprise the same fundamental components: neural activity fitting, BOLD-level fitting through the hemodynamic output layer, SC inference, and final refinement with FCD constraints. The complete optimization schedules, including trainable variables, output space, active loss terms, epoch counts, learning rates, batch sizes, warm-up periods, and training simulation durations, are summarized in Supplementary Table 2. The same schedule is used across the Glasser, Desikan–Killiany, and Harvard–Oxford human analyses.

### Computational implementation and empirical evaluation

The full optimization procedure is organized into training, validation, and test phases. Training is performed on GPUs using BPTT. Because matching the full empirical time-series length would require excessively long BPTT unrolls, increasing GPU memory burden and potentially compromising numerical precision, training instead uses multiple stochastic batches with relatively short simulation durations. Depending on the optimization stage and dataset, training uses batch sizes of 15–100 independent noise realizations, with each trajectory simulated for 1–3 min. This design provides stochastic robustness while keeping the BPTT graph computationally tractable.

Validation is performed on CPUs after saved training epochs become available. Because validation does not require gradient computation, it uses longer simulations matched to the corresponding empirical dataset duration, with one stochastic batch per candidate fit. For example, HCP validation simulations retain 14.4 min of BOLD activity after discarding the initial transient period. Validation simulations always include the hemodynamic output layer, regardless of whether the corresponding training stage uses neural activity-level or BOLD-level outputs. This places parameter states saved throughout the multistage curriculum in a common BOLD-level evaluation space. After each training stage is completed, its saved epochs are submitted for CPU-based validation while GPU training proceeds to the next step, thereby reducing the overall computational time.

During each optimization run, the parameter state at every epoch is saved. Each saved parameter state constitutes an epoch-specific candidate fit. Thus, the 2,500-epoch human optimization schedule shown in Supplementary Table 2 produces 2,500 candidate fits per optimization run. Candidate fits originating from the same run represent distinct parameter realizations sampled along a common optimization trajectory and are not described as separate optimization runs. For downstream inferred-SC analyses, eligibility is restricted to candidate fits saved after SC becomes trainable; earlier candidate fits remain part of the saved-epoch validation diagnostics but cannot enter inferred-SC selection or test analyses. For the fixed-SC controls in Fig. 3E–G, the EDR-derived connectome is held constant throughout optimization. All other trainable variables, output spaces, loss terms, optimizer settings, and epoch budgets match the corresponding inferred-SC schedule. Because SC never becomes trainable in these controls, all saved candidate fits are eligible for run-level validation selection. Each candidate fit is evaluated using a separately generated stochastic validation trajectory. Candidate fits are ranked using

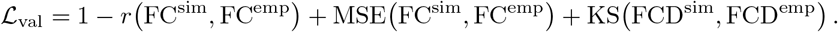

Within each optimization run, the 20 eligible candidate fits with the smallest ℒ_val_ proceed to validation-threshold screening. The same criterion is used for all loss-based fit ranking, including selection of the representative best fit and the validation-selected fit from each run. Building on the validation-based retention of multiple candidate parameter sets in pMFM [17], this procedure preserves multiple well-performing parameter states sampled along each optimization trajectory rather than reducing the trajectory to a single terminal estimate.

For both training and validation/test, simulated time series are downsampled to match the temporal resolution of the corresponding empirical fMRI data before FC and FCD evaluation. Unless otherwise stated, FC correlation and FC MSE are computed from the upper-triangular off-diagonal entries of the simulated and empirical FC matrices. For FCD-constrained training, FCD entries are computed from sampled 3-min time-series segments using 60-s sliding windows with a 10-s step, and the same sampling procedure is applied to the empirical data when computing the training loss. This approximation reduces GPU memory consumption and accelerates BPTT. In validation and test simulations, FCD is constructed from the complete post-warm-up trajectory using 60-s windows advanced by one downsampled time point; the same window and stride are used for empirical FCD. Empirical and simulated FCD entries are accumulated in 10,000 equal-width bins spanning [0, 1], and the corresponding normalized cumulative histograms define grid-based CDFs. The reported FCD KS distance is the maximum absolute difference between the empirical and simulated grid-based CDFs.

For the primary ensemble analyses, the 20 lowest-loss eligible candidate fits from each optimization run are screened using the three prespecified validation criteria for the corresponding dataset and parcellation:

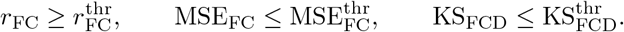

A candidate fit is designated a retained fit only if it belongs to the per-run top-20 set and satisfies all three criteria. The dataset-specific thresholds are reported in Supplementary Table 3. A single optimization run can therefore contribute multiple retained fits. For the empirical fit ensembles, we rank candidates and apply retention thresholds using validation metrics alone.

Every candidate fit entering a test-space analysis is evaluated using 30 independently generated stochastic test batches with the same simulation duration, sampling rate, hemodynamic output layer, and FC/FCD definitions used in validation. This test-evaluated set includes all retained fits, the run-level validation-selected fits used in run-level comparisons, and the 20 lowest-validation-loss eligible candidate fits from each optimization run displayed in Fig. 4E,F and Fig. 5E. For each evaluated fit, the 30 batch-specific FC matrices are averaged elementwise to obtain one test FC matrix, whereas FCD entries are pooled across the 30 batch-specific FCD matrices to obtain one test FCD distribution. All downstream test metrics and summaries are computed from these within-fit aggregates. Consequently, each evaluated fit contributes exactly one test FC estimate, one test FCD distribution, and one value to each derived test summary.

For comparisons designed to contribute one result per completed optimization run, the eligible candidate fit minimizing ℒ_val_ within each run is termed the validation-selected fit for that run. Within each dataset and model setting, the validation-selected fit with the smallest ℒ_val_ across completed runs is designated the representative best fit and is used only for the single-fit visualizations specified in the corresponding figure captions. Fig. 3B includes the validation-selected fit from every completed Glasser inferred-SC optimization run. The fixed-SC versus inferred-SC comparisons in Fig. 3E–G, the multi-stage ablation analysis in Supplementary Fig. 1, and the benchmark comparison in Supplementary Fig. 4 likewise use run-level validation-selected fits without applying the downstream retention thresholds.

An optimization run contributing at least one retained fit is termed a retained run. For the crossrun consistency analysis in Supplementary Fig. 3, each retained run contributes only its retained fit with the smallest ℒ_val_. The test-performance distributions in Fig. 4E,F and Fig. 5E include all 20 lowest-validation-loss eligible candidate fits from every optimization run, whether or not they satisfy the validation thresholds; the red lines indicate the least favorable test performance among the threshold-qualified retained fits. All other fit distributions and across-fit regional-parameter or SC summaries use retained fits unless otherwise stated. When regional parameters or SC are displayed as ensemble averages, retained fit-level estimates are aggregated elementwise after applying the same preprocessing and normalization used for each individual fit.

### RSFC hierarchy construction

RSFC hierarchy measures are derived using diffusion embedding of empirical FC, following Margulies et al. [32]. For the Glasser and Desikan–Killiany analyses, an atlas-specific principal gradient is derived from the corresponding empirical FC target. For the Harvard–Oxford analysis, a single reference FC matrix is formed by elementwise averaging the empirical NC and MDD group-average FC targets,

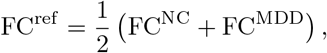

and one principal gradient derived from this reference matrix is applied unchanged to both the NC and MDD fit ensembles. Diffusion embedding provides a nonlinear low-dimensional representation of similarities among regional connectivity profiles, and the leading embedding component is used as the RSFC hierarchy measure. Because the polarity of an embedding component is mathematically arbitrary, each gradient is oriented such that higher values correspond to the transmodal/default-mode end of the principal gradient.

### SC–tracer alignment analysis

We compare inferred coupling with tracer weights in the macaque Markov and marmoset Paxinos parcellations. For each injection target, FLN is the number of labeled neurons in a given source area divided by the total number of labeled neurons outside the target in the same hemisphere. Diffusion-MRI-derived SC and the EDR initialization are analyzed using the same procedure.

Although StaGIC infers a directed connectivity matrix, diffusion tractography does not resolve projection direction. To place all candidate structural estimates on the same representational footing for this benchmark, StaGIC-inferred, dMRI-derived, and EDR-derived SC matrices are symmetrized before comparison,

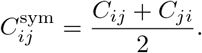

Tracer-derived FLN values are retained in their original directed, target-normalized representation because tracer coverage is incomplete across target columns and reciprocal averaging would introduce coverage-dependent distortions. This symmetrization is applied only for the cross-modal anatomical benchmark and does not alter the directed SC used during model fitting or in other analyses.

For each trainable-SC method, the symmetrized SC matrices are first averaged elementwise across the fit ensemble used in the corresponding analysis, and the distance-resolved Spearman correlations are then computed from this ensemble-mean SC. The main cross-species analysis uses retained fits, whereas the benchmark analysis in Supplementary Fig. 4 uses all validation-selected fits from completed optimizations.

Comparisons are restricted to tracer-covered off-diagonal ROI pairs, including both positive and measured zero-valued FLN entries; ROI pairs without tracer coverage are excluded. The same tracer-defined support is applied to StaGIC-inferred, dMRI-derived, and EDR-derived SC and does not depend on the value or sparsity pattern of any candidate SC matrix. Spearman’s rank correlation is used because FLN spans several orders of magnitude [25, 27]: on the raw scale, Pearson correlation would be disproportionately driven by the strongest projections, whereas logarithmic transformation would require either excluding measured zeros or assigning an arbitrary pseudocount. Rank correlation limits the leverage of the strongest edges while preserving their ordinal organization and retaining measured zero-valued FLN entries as tied ranks. Because the same calculation is repeated from the full tracer-covered edge set to progressively longer-distance subsets, this metric supports both whole-range and long-range comparisons without changing the treatment of zero-valued projections. For each minimum distance threshold *d*, ROI pairs with inter-regional distance greater than *d* are retained, and Spearman’s rank correlation is computed between each symmetrized SC estimate and the measured FLN values across the retained edges. No pseudocount or logarithmic transformation is applied.

For each species, we restrict the analysis to distance thresholds that retain at least 25% of the ROI pairs included at *d* = 0. The largest thresholds are 25.38 mm in macaque and 14.38 mm in marmoset, both above the corresponding 5*λ* cutoff.

### MDD latent-circuit contrast analysis

Latent SC contrasts are evaluated over the Cartesian product of the retained NC and MDD fit ensembles, without one-to-one matching between fits. For these analyses, inferred long-range coupling is represented by

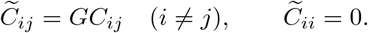

Let 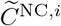 and 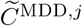 denote the effective coupling matrices from retained NC fit *i* and MDD fit *j*, respectively. All combinations of NC and MDD fits are evaluated.

For each cross-group fit pair (*i, j*) and ROI *k*, the signed efferent and afferent differences in effective inferred coupling are defined as

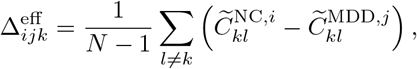

and

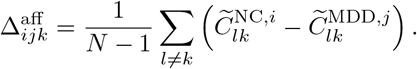

Positive values therefore indicate stronger inferred coupling in NC than in MDD. For Fig. 6C, ROI-level values are averaged separately within the MDD-annotated and other ROI sets for each NC–MDD fit pair, and the resulting cross-fit-pair distributions are displayed descriptively.

Projection-profile dissimilarity is evaluated separately for efferent and afferent connectivity. For each ROI and NC–MDD fit pair, self-connections are excluded and projection-profile dissimilarity is defined as one minus the Pearson correlation between the corresponding NC and MDD projection profiles. ROI-level dissimilarities are then averaged within the MDD-annotated and other ROI sets for each fit pair, yielding the cross-fit-pair distributions shown in Fig. 6E.

For Fig. 6B, ROI-level signed differences in effective inferred coupling are averaged across all NC–MDD cross-fit pairs before cortical-surface visualization. For Fig. 6D, each retained fit contributes one absolute Pearson correlation between the corresponding regional parameter vector and the RSFC hierarchy gradient.

### Cross-run consistency analysis

Cross-run consistency is evaluated across retained optimization runs from the primary Glasser analysis. An optimization run is retained if it contributes at least one retained fit, and each retained run is represented by its retained fit with the smallest ℒ_val_. Thus, no optimization run contributes more than one parameter realization to this analysis. For *w*_*i*_, *I*_*i*_, and *σ*_*i*_, pairwise similarity between run-level fits is quantified by Pearson correlation across ROIs. Regional-parameter and SC comparisons use the selected retained parameter states, whereas FC and FCD comparisons use their corresponding 30-batch aggregated test outputs defined above. SC similarity is computed across all off-diagonal directed entries, whereas FC similarity is computed across upper-triangular off-diagonal entries. Parameter, SC, and FC dissimilarity is reported as 1 − *r*. FCD dissimilarity is quantified by the KS distance between FCD-entry distributions. For the summary comparison, consistency is expressed as *r* for *w*_*i*_, *I*_*i*_, *σ*_*i*_, SC, and FC, and as 1−KS for FCD.

### Multistage ablation and synthetic parameter recovery

To evaluate the contribution of the optimization curriculum, the original StaGIC schedule is compared with three ablation strategies: all parameters trainable from the beginning, all loss terms active from the beginning, and both all parameters and all loss terms active from the beginning. All strategies use the same model formulation, initialization, total optimization budget, and evaluation procedure, such that their primary difference is the ordering of parameter and loss activation.

Synthetic recovery is evaluated using three distinct well-fitting Glasser parameterizations as ground-truth generators. Candidate parameterizations are first restricted to models with test FC correlation greater than 0.88 and FCD KS distance less than 0.24. To avoid selecting nearly identical latent solutions, three ground-truth models are then chosen using a maximin diversity criterion based jointly on regional parameters, effective SC, FC, and FCD. Pairwise MSE and 1 − *r* are computed for *w, I, σ*, effective SC, and FC, together with the KS distance between FCD cumulative distributions. Each of the 11 distance matrices is min–max normalized across candidate models; the normalized distances are averaged with unit weight for all measures except FCD KS distance, which is assigned weight 2. This procedure yields three functionally well-fitting but distinct latent parameter configurations.

For each ground-truth configuration, synthetic BOLD data are generated using the same Glasser forward model and hemodynamic readout as in the main analysis, with a 2-min warm-up followed by 14.4 min of retained BOLD activity. Synthetic observations and all subsequent optimization runs use mutually independent stochastic noise realizations, so that optimization never reuses the noise trajectory that generated the target data. The resulting synthetic FC and FCD statistics are used as fitting targets. Each optimization strategy is evaluated with 10 independent optimizations for each of the three ground-truth configurations, yielding 30 independent optimizations per strategy. All strategies use the same initialization as the standard Glasser analysis, including the same EDR-based SC initialization. Functional reconstruction is quantified by FC correlation, FC MSE, and FCD KS distance. Recovery of *w*_*i*_, *I*_*i*_, *σ*_*i*_, and the effective SC matrix 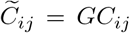 for *i* ≠ *j*, with 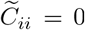, is quantified by Pearson correlation and MSE relative to the known ground-truth values.

### Statistical analysis

All hypothesis tests are two-sided. The Mann–Whitney U test is used for comparisons between participant-level observations or fit-level test summaries, as specified below.

The fixed-SC versus inferred-SC comparisons in Fig. 3E–G are presented descriptively, with each completed independent optimization run contributing one validation-selected fit. Bars and confidence intervals summarize variation across these run-level computational replicates; no hypothesis tests are performed for these comparisons. The runs do not represent independent biological replicates. Benchmark comparisons between StaGIC and pMFM or TVB-Optim in Supplementary Fig. 4 are likewise presented descriptively, and no hypothesis tests are performed.

For the MDD functional summaries in Fig. 5G,H, empirical NC–MDD comparisons use individual participants as independent biological observations, with *n* = 968 for NC and *n* = 1,150 for MDD. Simulated comparisons use retained fit-level test summaries, with *n* = 288 for NC and *n* = 246 for MDD. Each retained fit is evaluated using its own independently generated test-noise ensemble, the resulting test batches are aggregated within fit, and that fit contributes exactly one simulated summary. All empirical and simulated summaries are computed from the same upper-triangular off-diagonal FC entries.

Retained fits originating from the same optimization run are distinct validation-qualified parameter realizations sampled along a common optimization trajectory and are not interpreted as independent optimization runs. Conditional on these retained parameter states, their test summaries are generated from non-overlapping stochastic test-noise ensembles. Accordingly, tests across simulated fit-level summaries quantify differences between the conditional predictive ensembles generated by the NC and MDD group models. These tests do not constitute subject-level biological inference or an estimate of variability across fully independent optimization runs.

For the MDD analyses, hypothesis tests are restricted to global functional connectivity and global functional differentiation in Fig. 5G,H, quantified respectively by the mean and standard deviation of upper-triangular off-diagonal FC entries. The latent regional-parameter and SC contrasts shown in Fig. 6 are presented descriptively, and no hypothesis tests are performed for these panels. These latent analyses are intended to generate candidate group-level circuit hypotheses and should not be interpreted as subject-level statistical inference.

Pearson correlation is used for FC, regional-parameter, hierarchy, and cross-run correspondence where indicated. SC–tracer correspondence is quantified using Spearman’s rank correlation, retaining measured zero-valued tracer entries and applying the corresponding anatomical-distance threshold. FCD similarity is evaluated using the KS distance between FCD-entry distributions. For the cross-run consistency analysis, parameter, SC, and FC dissimilarity is defined as 1 − *r*, whereas FCD dissimilarity is quantified directly by the KS distance.

No correction for multiple comparisons is applied, and all reported *P* values are therefore uncorrected. Statistical significance is defined as *P <* 0.05. Significance annotations are denoted as ns, *P* ≥ 0.05; *, *P <* 0.05; **, *P <* 0.01; ***, *P <* 0.001; and ****, *P <* 0.0001.

Unless otherwise specified, summary bars represent the mean across the corresponding observations. All error bars denote 95% confidence-interval half-widths around the corresponding mean. Unless otherwise stated, confidence intervals are computed using the two-sided Student’s *t* distribution:

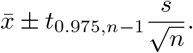

For participant-level summaries, *n* denotes the number of independent participants. For summaries across retained fits, *n* is the number of fits. Each fit contributes one FC statistic computed from its mean test FC matrix in Fig. 5G,H, or one correlation per regional parameter computed directly from the fitted map in Fig. 3D and Fig. 6D. The reported intervals summarize fit-level means for the corresponding group target. For run-level summaries, *n* denotes the number of completed optimization runs, with each run contributing one validation-selected fit. For empirical MDD summaries in Fig. 5G,H, confidence intervals are computed across participants, whereas simulated intervals are computed across retained fit-level test summaries. For Fig. 6C,E, all NC–MDD cross-fit-pair values are displayed descriptively. Because different fit pairs share individual NC or MDD fits, these pairwise values are not used for confidence-interval estimation or hypothesis testing. For Fig. 6D, each retained fit contributes one parameter–hierarchy correlation, and the displayed interval is computed across these fit-level values.

### Comparison with pMFM and TVB-Optim

For comparison with pMFM, fixed-SC models are evaluated under the Desikan and Glasser parcellations using FC correlation and FC MSE. For comparison with TVB-Optim, the available released implementation is evaluated in the inferred-SC setting under the human Desikan and Glasser parcellations and in macaque and marmoset datasets. Non-human primate comparisons additionally evaluate the correspondence between inferred SC and tracer-derived connectivity across anatomical distance thresholds. SC–tracer correspondence is quantified using Spearman’s rank correlation with zero-valued tracer entries retained. The resulting functional and anatomical metrics are evaluated using the same definitions as in the corresponding StaGIC analyses.

For each benchmark, the same empirical FC targets and structural inputs are used across methods wherever supported by the released implementation. Each method is run with 20 independent optimizations under the corresponding published or released parameterization. Model outputs are evaluated using the same downstream FC and SC–tracer metric implementations used for StaGIC. The pMFM benchmark uses the Standalone Kong2021 pMFM v0.21.1 release (https://github.com/ThomasYeoLab/StandaloneKong2021pMFM/releases/tag/v0.21.1), and the TVB-Optim benchmark uses the tvboptim v0.4.0 release (https://github.com/virtual-twin/tvboptim/releases/tag/v0.4.0).

## Software implementation

The main software dependencies are Python 3.9, JAX 0.4 [58], and BrainPy 2.6 [59].

## Data availability

The source datasets analyzed in this study are available from the WU–Minn Human Connectome Project, REST-meta-MDD, and the macaque, marmoset, and tracer resources cited in Methods, subject to the access conditions of the originating repositories. No new physical materials were generated.

## Code availability

The core StaGIC method code, including the complete training architecture and a runnable HCP Human Glasser example with all derived inputs required to execute it, is openly available at GitHub (https://github.com/BrianYahoo/StaGIC).

## Acknowledgements

This work was supported by National Key R&D Program of China 2025YFA1016700 (D.Z., S.Li); Shanghai Municipal Commission of Science and Technology with Grant No. 24JS2810400 (D.Z., S.Li); National Natural Science Foundation of China Grant 12631018, 12271361 (S.Li); National Natural Science Foundation of China with Grant No. 92570202, 12225109 (D.Z.); and the Student Innovation Center at Shanghai Jiao Tong University. Data were provided in part by the members of REST-meta-MDD Consortium.

## Author contributions

Conceptualization: X.C., D.Z., and S.L., Methodology: B.Y., Z.J., X.C. Validation: B.Y. Formal analysis: B.Y. Investigation: B.Y. Data curation: B.Y. Visualization: B.Y. Project administration: X.C., D.Z., and S.L. Writing–original draft: B.Y. Writing–review and editing: B.Y., Z.J., X.C., D.Z., and S.L. Supervision: X.C., D.Z. and S.L. All authors reviewed and approved the final manuscript.

## Additional information

Supplementary Information accompanies this manuscript and contains Supplementary Tables 1–3 and Supplementary Figs. 1–5. Correspondence and requests for materials should be addressed to Xiaoyu Chen, Douglas Zhou or Songting Li.

