## Supplementary for "Multistage gradient-based whole-cortex modeling for joint inference of regional circuit heterogeneity and interregional structural connectivity"

### Supplementary Notes

The supplementary tables and figures provide literature-based MDD region annotations, dataset-specific optimization schedules and validation-retention criteria, hemodynamic implementation checks, multi-stage optimization trajectories and ablations, cross-run consistency analyses, and direct comparisons with pMFM and TVB-Optim in functional fitting and connectome inference.

**Supplementary Table 1  Literature-informed mapping of MDD-annotated cortical regions to the Harvard–Oxford atlas.** Candidate cortical systems were identified from meta-analyses and large multisite studies of MDD and manually mapped to the closest Harvard–Oxford labels listed here [1–3]. The cited evidence supports involvement of the broader region or system and not necessarily the exact atlas subdivision. All remaining cortical ROIs form the descriptive comparison set ("other ROIs").

| ROI Name | MDD-annotated | Reference |
|---|---|---|
| Insular Cortex | Yes | [1, 2] |
| Middle Frontal Gyrus | Yes | [1, 2] |
| Inferior Frontal Gyrus pars triangularis | Yes | [3, 4] |
| Inferior Frontal Gyrus pars opercularis | Yes | [3, 4] |
| Temporal Pole | Yes | [5] |
| Superior Temporal Gyrus anterior division | Yes | [5, 6] |
| Middle Temporal Gyrus posterior division | Yes | [3, 6] |
| Frontal Medial Cortex | Yes | [1, 7] |
| Subcallosal Cortex | Yes | [8] |
| Paracingulate Gyrus | Yes | [2] |
| Cingulate Gyrus anterior division | Yes | [2, 7] |
| Cingulate Gyrus posterior division | Yes | [1, 7] |
| Frontal Orbital Cortex | Yes | [2, 9] |
| Parahippocampal Gyrus anterior division | Yes | [3, 4] |
| Parahippocampal Gyrus posterior division | Yes | [3, 4] |
| Temporal Fusiform Cortex anterior division | Yes | [2] |

**Supplementary Table 2 StaGIC optimization schedules for SC inference across species.** The same schedule is used across all human datasets and parcellations, whereas macaque and marmoset use species-specific schedules. LR denotes learning rate, and Batch denotes the number of independent stochastic noise trajectories simulated in parallel.

| Species | Step | Trainable variables | Output | Loss | Epochs | LR | Batch | Warm-up | Simulation |
|---|---|---|---|---|---|---|---|---|---|
| Human | 1 | $G, w, I, \sigma$ | Activity | FC | 1000 | $10^{-2}$ | 100 | 10 s | 60 s |
| | 2 | SC,$w, I, \sigma$ | Activity | FC | 500 | $10^{-3}$ | 100 | 10 s | 60 s |
| | 3 | SC,$w, I, \sigma$ | Hemodynamic | FC | 500 | $10^{-4}$ | 25 | 30 s | 180 s |
| | 4 | SC,$w, I, \sigma$ | Hemodynamic | FC+FCD | 500 | $10^{-5}$ | 15 | 30 s | 180 s |
| Macaque | 1 | $G, w, I, \sigma$ | Activity | FC | 1000 | $10^{-2}$ | 100 | 10 s | 60 s |
| | 2 | $G, w, I, \sigma$ | Activity | FC | 300 | $10^{-3}$ | 100 | 10 s | 60 s |
| | 3 | $G, w, I, \sigma$ | Hemodynamic | FC | 200 | $10^{-4}$ | 25 | 30 s | 180 s |
| | 4 | SC,$w, I, \sigma$ | Hemodynamic | FC | 500 | $10^{-4}$ | 25 | 30 s | 180 s |
| | 5 | SC,$w, I, \sigma$ | Hemodynamic | FC+FCD | 500 | $10^{-3}$ | 15 | 30 s | 180 s |
| | 6 | SC,$w, I, \sigma$ | Hemodynamic | FC+FCD | 500 | $10^{-4}$ | 15 | 30 s | 180 s |
| Marmoset | 1 | $G, w, I, \sigma$ | Activity | FC | 1000 | $10^{-2}$ | 100 | 10 s | 60 s |
| | 2 | $G, w, I, \sigma$ | Activity | FC | 300 | $10^{-3}$ | 100 | 10 s | 60 s |
| | 3 | $G, w, I, \sigma$ | Hemodynamic | FC | 200 | $10^{-4}$ | 25 | 30 s | 180 s |
| | 4 | SC,$w, I, \sigma$ | Hemodynamic | FC | 500 | $10^{-4}$ | 25 | 30 s | 180 s |
| | 5 | SC,$w, I, \sigma$ | Hemodynamic | FC+FCD | 500 | $10^{-5}$ | 15 | 30 s | 180 s |
| | 6 | SC,$w, I, \sigma$ | Hemodynamic | FC+FCD | 500 | $10^{-5}$ | 15 | 30 s | 180 s |

**Supplementary Table 3 Dataset- and parcellation-specific validation criteria for downstream inferred-SC ensemble analyses.** Each saved parameter state constitutes a candidate fit. After inferred-SC eligibility is restricted to epochs saved after SC becomes trainable, the 20 eligible candidate fits with the lowest combined validation criterion are selected from each optimization run. Candidate fits within this per-run top-20 set that satisfy all three validation thresholds are designated retained fits. A single optimization run can therefore contribute multiple retained fits. All ranking and threshold criteria are applied exclusively to validation metrics.

| Dataset / parcellation | FC corr $\geq$ | FC MSE $\leq$ | FCD KS $\leq$ | Retained fits |
|---|---|---|---|---|
| Human / Glasser | 0.88 | 0.03 | 0.2 | 373 |
| Human / Desikan | 0.90 | 0.02 | 0.1 | 252 |
| NC / Harvard–Oxford | 0.85 | 0.02 | 0.3 | 288 |
| MDD / Harvard–Oxford | 0.85 | 0.02 | 0.3 | 246 |
| Macaque / Markov | 0.86 | 0.02 | 0.25 | 319 |
| Marmoset / Paxinos | 0.83 | 0.03 | 0.3 | 209 |

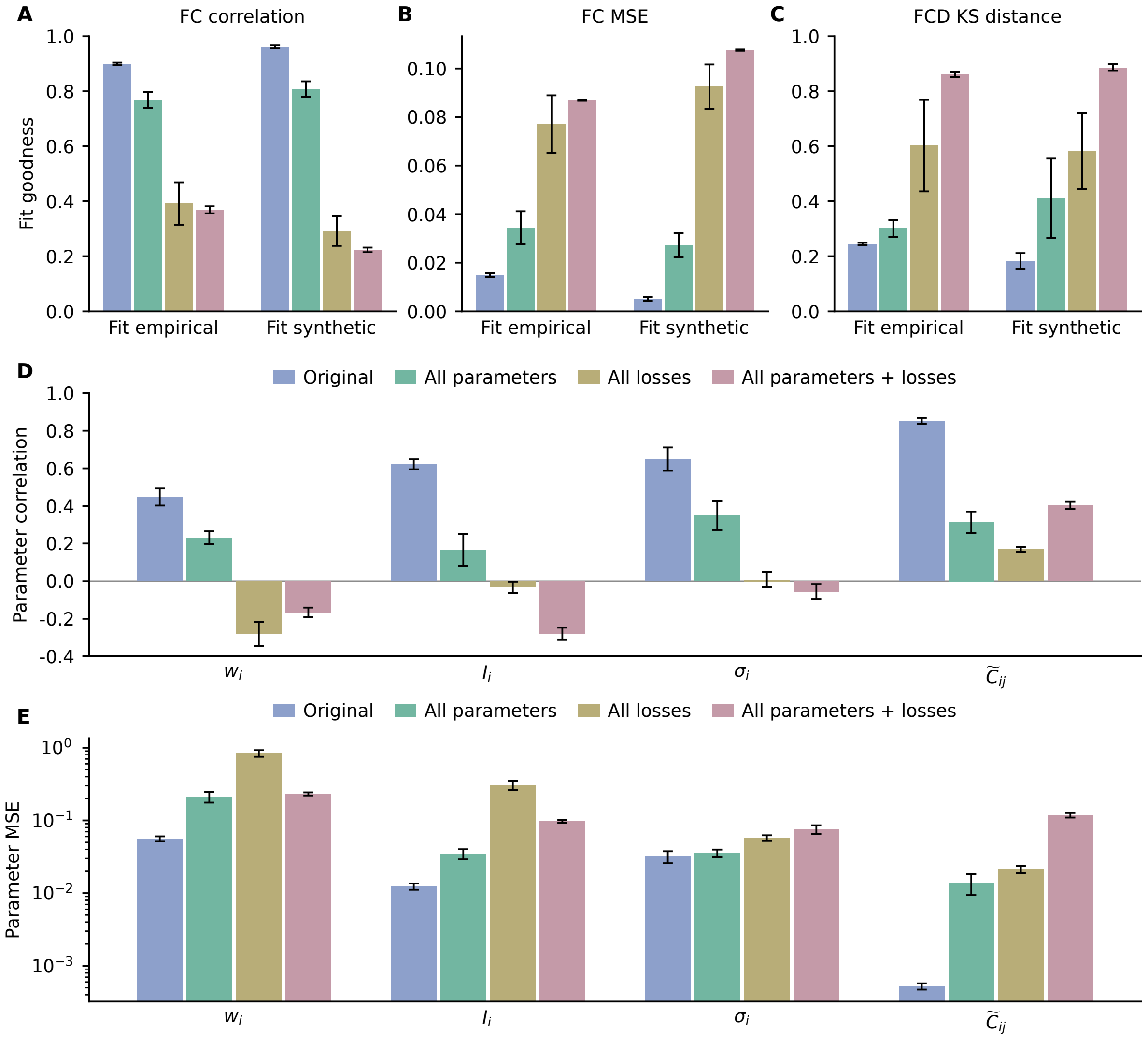


**Supplementary Fig. 1 Multistage optimization improves functional fitting and parameter recovery.** Four strategies are compared: Original StaGIC; All parameters, with all parameters trainable from the beginning; All losses, with all loss terms introduced from the beginning; and All parameters + losses, with both activated from the beginning. (**A**–**C**) FC correlation, FC MSE, and FCD KS distance when fitting empirical Glasser data or synthetic data. (**D**, **E**) Correlation and MSE between recovered and ground-truth $w_i$, $I_i$, $\sigma_i$, and effective SC $\widetilde{C}_{ij}$ for the synthetic-data experiment. Bars and error bars indicate mean $\pm$ 95% Student's $t$ CI half-width across 30 independent optimizations per strategy.

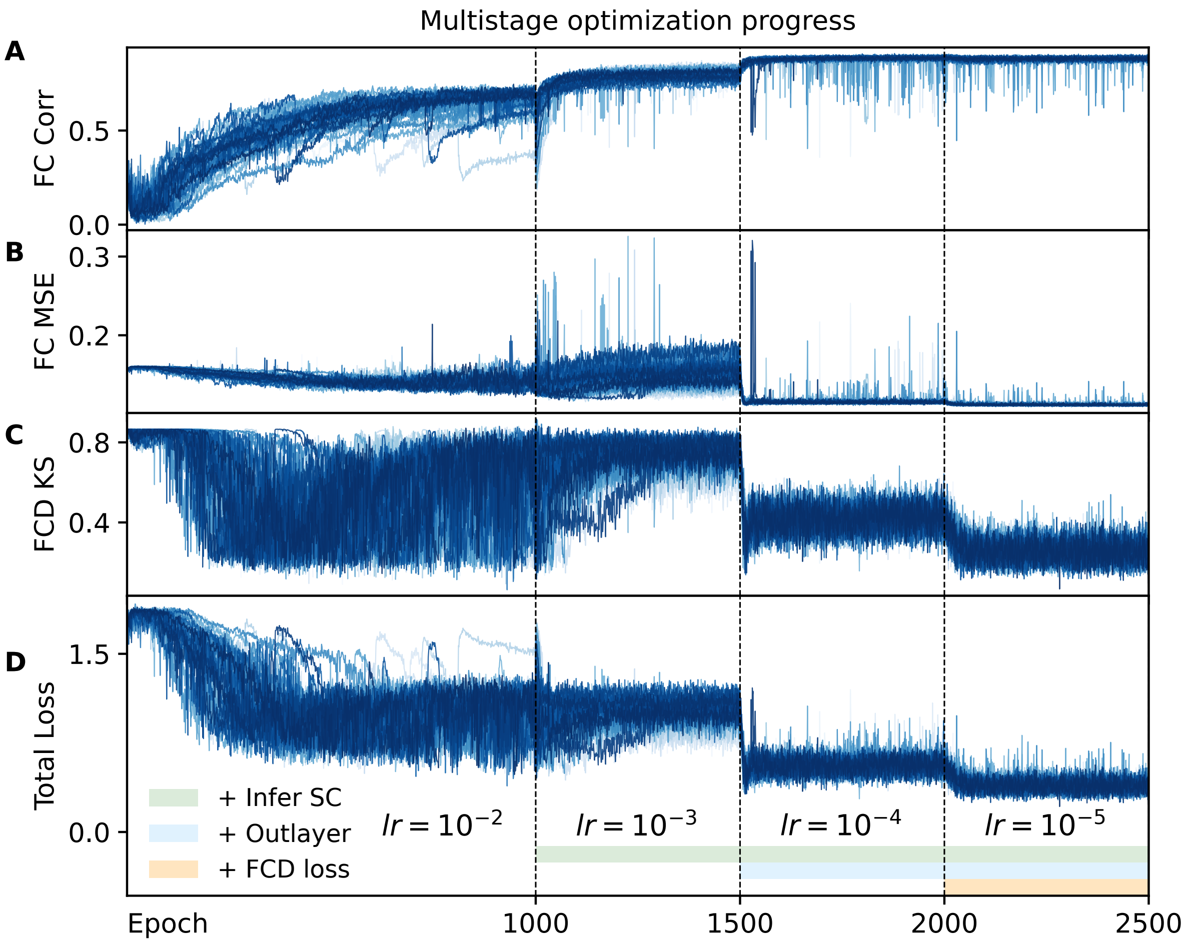


**Supplementary Fig. 2 Validation trajectories across the multistage Glasser analysis.** The four panels show validation metrics evaluated at saved epochs for (**A**) FC correlation, (**B**) FC MSE, (**C**) FCD KS distance, and (**D**) the combined validation criterion $\mathcal{L}_{\text{val}} = 1 - r_{\text{FC}} + \text{MSE}_{\text{FC}} + \text{KS}_{\text{FCD}}$ used to rank saved-epoch candidate fits. Each curve traces the candidate fits sampled from one complete optimization run. Vertical dashed lines mark the boundaries of the multistage curriculum, and colored bars indicate the stages in which SC, FCD loss, and the BOLD output layer are active.

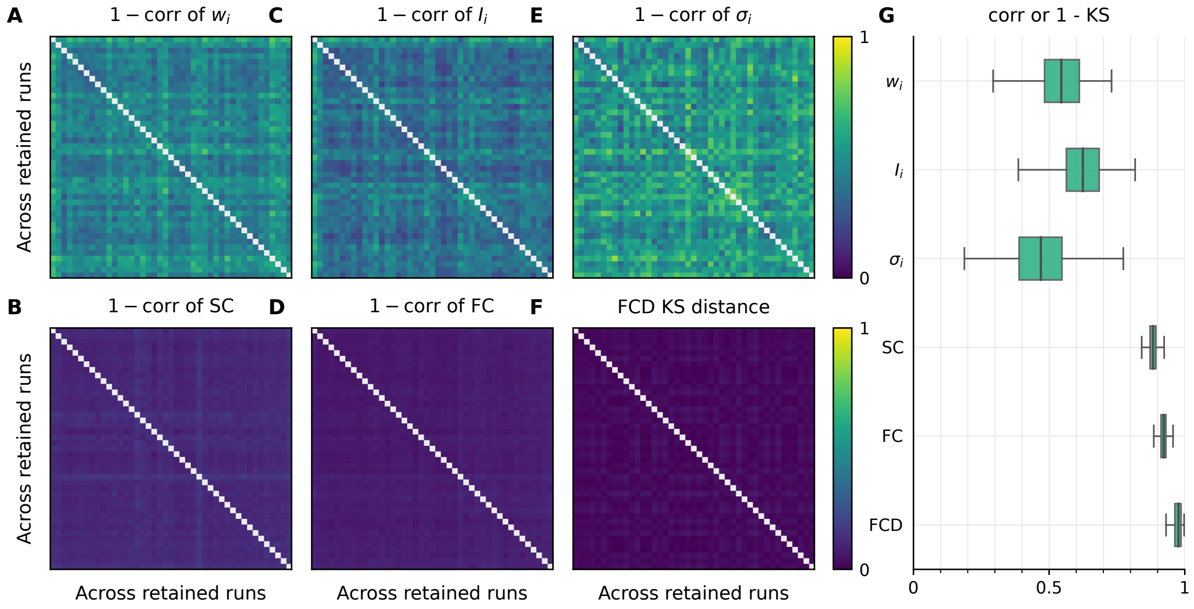


**Supplementary Fig. 3 Cross-run consistency reveals a stable component of inferred SC.** (**A–F**) Pairwise dissimilarity for $w_i$, SC, $I_i$, FC, $\sigma_i$, and FCD, respectively, across optimization runs contributing at least one retained fit. Each run is represented by its retained fit with the lowest validation loss. Dissimilarity is quantified by $1 - r$ for parameters, SC, and FC, and by the KS distance for FCD. (**G**) Distributions of pairwise cross-run consistency, shown as Pearson correlation for $w_i$, $I_i$, $\sigma_i$, SC, and FC and as $1 - \text{KS}$ for FCD; boxes show medians and interquartile ranges.

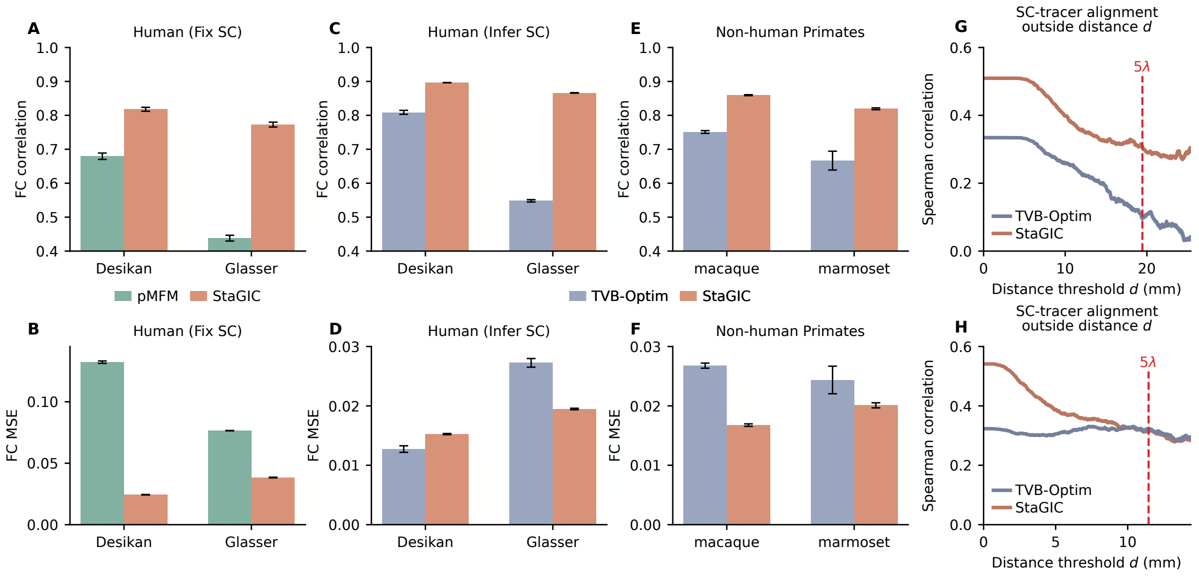


**Supplementary Fig. 4 Comparison of StaGIC with pMFM and TVB-Optim in functional fitting and connectome inference.** (**A**, **B**) Human fixed-SC comparison between pMFM and StaGIC under the Desikan and Glasser parcellations, evaluated by FC correlation and FC MSE. (**C**, **D**) Human inferred-SC comparison between TVB-Optim and StaGIC under the Desikan and Glasser parcellations. (**E**, **F**) FC-fitting comparison between TVB-Optim and StaGIC for macaque and marmoset. (**G**, **H**) Distance-dependent SC–tracer alignment for macaque and marmoset, quantified using Spearman's rank correlation over tracer-covered ROI pairs with measured zero-valued entries retained; red dashed lines indicate the species-specific $5\lambda$ threshold. For StaGIC and TVB-Optim, SC–tracer curves are computed from the elementwise mean SC across all validation-selected fits before correlation. Bars and error bars in (**A**)–(**F**) indicate mean ± 95% Student's $t$ CI half-width across all validation-selected fits from completed independent optimizations.

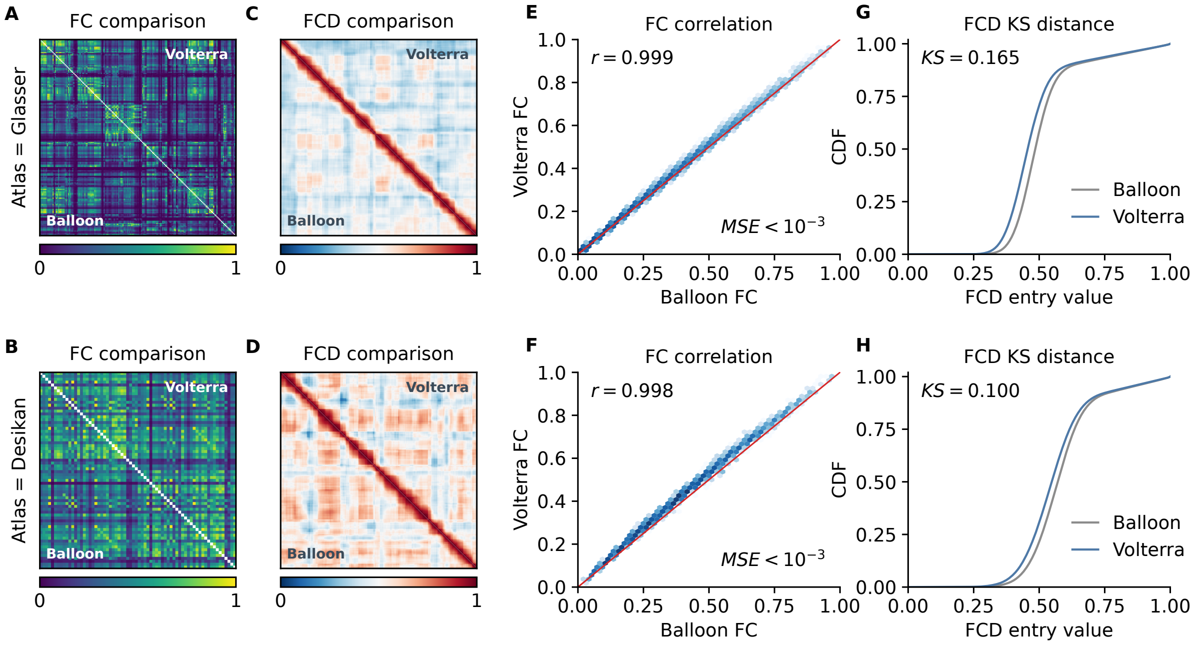


**Supplementary Fig. 5 Comparison between approximate and reference model implementations.** (**A**, **B**) FC matrix comparison for the representative best fits under the Glasser and Desikan parcellations. In each matrix, the lower triangle shows Balloon–Windkessel FC and the upper triangle shows Volterra FC. (**C**, **D**) FCD matrix comparison for the same representative best fits; lower triangles show Balloon–Windkessel FCD and upper triangles show Volterra FCD. (**E**, **F**) FC-entry scatter plots comparing Balloon–Windkessel and Volterra outputs. (**G**, **H**) FCD-entry CDF comparisons between Balloon–Windkessel and Volterra outputs. The Volterra implementation uses the Softplus neural transfer function, and the Balloon–Windkessel implementation uses the original Abbott–Chance transfer function. Both implementations use the same fitted neural parameters for each representative best fit; panels (**C**) and (**D**) show the first prespecified test realization. Because empirical FCD entries are pooled across runs before constructing the cumulative distribution, no single empirical FCD matrix is defined for comparison in panels (**C**) and (**D**).
